# The neuropeptide Galanin is required for homeostatic rebound sleep following increased neuronal activity

**DOI:** 10.1101/479634

**Authors:** Sabine Reichert, Oriol Pavón Arocas, Jason Rihel

## Abstract

Sleep pressure homeostatically increases during wake and dissipates during sleep, but the molecular signals and neuronal substrates that measure homeostatic sleep pressure remain poorly understood. We present a pharmacological assay in larval zebrafish that generates acute, short-term increases in wakefulness followed by sustained rebound sleep after washout. The intensity of global neuronal activity during drug-induced wakefulness predicted the amount of subsequent rebound sleep. Whole brain mapping with the neuronal activity marker phosphorylated extracellular signal–regulated kinase (pERK) identified preoptic Galanin-expressing neurons as selectively active during rebound sleep, and the relative induction of *galanin* transcripts was predictive of total rebound sleep time. Galanin is required for sleep homeostasis, as *galanin* mutants almost completely lacked rebound sleep following both pharmacologically induced neuronal activity and physical sleep deprivation. These results suggest that Galanin plays a key role in responding to sleep pressure signals derived from neuronal activity and functions as an output arm of the vertebrate sleep homeostat. (word count: 158).

## Introduction

Animals sleep at specific times of day. This timing, as well as the depth and quantity of sleep, are regulated by two interacting processes: a circadian, 24-hour process that sets sleep relative to the phase of the day/night cycle and a homeostatic process that increases the drive to sleep as a function of prior wakefulness (Borbely, 1982). While the molecular and neuronal underpinnings of circadian rhythms are fairly well understood, the mechanisms that measure and actuate homeostatic sleep pressure are only starting to be elucidated in flies (Donlea, 2017; Liu et al., 2016) and in vertebrates (Funato et al., 2016).

Because extended wakefulness is followed by longer and more intense rebound sleep (Borbely and Achermann, 1999), many hypotheses of sleep homeostasis converge on metabolic or synaptic correlates of wake-driven neuronal activity. For example, hypotheses of neuronal fatigue suggest that during extended wake, neurons can no longer sustain firing rates after the depletion of critical resources, such as glycogen or lipids essential for secretory vesicles (Benington and Heller, 1995; Scharf et al., 2008; Vyazovskiy and Harris, 2013). Active neurons may also produce sleep-signalling substances that are direct by-products of metabolic activity, such as extracellular adenosine (Porkka-Heiskanen and Kalinchuk, 2011), intracellular calcium (Ode et al., 2017), or nitric oxide (Kalinchuk et al., 2010). Alternatively, it has been proposed that activity-dependent strengthening of synaptic potentiation during wake may enhance synaptic coupling and neuronal synchrony, which in turn is associated with synaptic downscaling and neuronal plasticity during sleep (Tononi and Cirelli, 2016). However, it is still unclear how these correlates, alone or in combination, relate to the poorly understood mechanisms orchestrating sleep homeostasis.

A critical question to be addressed is how these still ill-defined correlates of wake-driven neuronal activity are sensed at the level of single neurons and transformed into global, whole organism sleep. One of the major hallmarks of rebound sleep following sleep deprivation in mammals is an increase in cortical slow-wave activity (SWA) during non-REM (NREM) sleep, which dissipates in intensity as sleep progresses (Bellesi et al., 2014). However, SWA intensity is not uniform across the brain but is locally increased during NREM sleep as a function of local brain use during prior wakefulness (Finelli et al., 2001; Huber et al., 2004; Werth et al., 1997). These observations have led to the concept of local “sleep”, in which homeostatic sleep pressure is predominantly sensed and regulated by assemblies of active neurons (Krueger et al., 2008). In this model, wake-active cortical neurons increase sleep pressure locally, either through the release of sleep-promoting substances or via changes in synaptic strength, which alters their firing properties to drive locally increased SWA (Roy et al., 2008). Less clearly explained by theories of local sleep is how local increases of SWA in neuronal assemblies could co-ordinately drive whole organismal sleep, although mathematical modelling has suggested whole organism sleep is an emergent property of the number of simultaneous “offline” assemblies (Roy et al., 2008). However, behavioral sleep is maintained even after local increases in sleep pressure have dissipated and local, wake-like neuronal activity patterns have returned, (Finelli et al., 2001; Werth et al., 1997), suggesting that non-local factors contribute to reinforce and maintain appropriate whole organismal sleep.

An alternative view hypothesizes that sleep is imposed upon the brain by the activity of centrally acting, circadian- and/or homeostatic-sensitive sleep-promoting neurons. Numerous classes of neuronal subpopulations in both Drosophila and mammals have been identified that are capable of driving sleep and wake when stimulated by opto- and chemogenetic approaches (Chen et al., 2018; Chung et al., 2017; Donlea et al., 2014; Donlea et al., 2018; Kroeger et al., 2018; Seidner et al., 2015; Xu et al., 2015). In addition, focal lesions of some brain nuclei, for example the mammalian ventro-lateral preoptic area (VLPO), can cause severe insomnia (Lu et al., 2000), demonstrating that a loss of a small number of critical neurons can have a large effect on sleep. Several of these sleep-inducing neurons are either directly or indirectly sensitive to homeostatic sleep signals. For example, in Drosophila, two classes of sleep regulatory neurons change electrophysiological properties in response to sleep need (Donlea, 2017; Liu et al., 2016). In vertebrates, neurons in both the VLPO and median preoptic nucleus increase firing in response to sleep pressure following sleep deprivation (Alam et al., 2014; Szymusiak et al., 1998; Szymusiak and McGinty, 2008). However, while sleep-promoting VLPO neurons are sensitive to manipulations of putative homeostatic sleep-pressure signals such as adenosine (Chamberlin et al., 2003; Kumar et al., 2013), the build-up of adenosine as a function of sleep loss has only been demonstrated in basal forebrain and cortex, not in the hypothalamus (Kalinchuk et al., 2010; Leenaars et al., 2018; Porkka-Heiskanen et al., 2000). In addition, the handful of genes that have been implicated in modulating sleep homeostasis, including *prion* (Tobler et al., 1997), the kinase *sik3* (Funato et al., 2016), and the circadian genes *clock*, *per3*, and *reverbα* (Franken and Dijk, 2009), are expressed across the brain and body. These observations raise the question whether signalling processes acting within specific sleep-regulatory neurons are critical for sensing and driving sleep homeostasis or whether sleep homeostasis signals are more globally distributed.

Here we develop a pharmacological assay that allows us to decouple sleep pressure from prior waking time and use it to show that pharmacologically induced neuronal activity drives homeostatic sleep pressure in zebrafish larvae. We find that following both drug- and physically-induced rebound sleep, transcription of the inhibitory neuropeptide *galanin* is upregulated in specific neurons of the preoptic area and hypothalamus as a function of prior neuronal activity. Furthermore, our genetic lesioning data shows that *galanin* is required for homeostatic rebound sleep, demonstrating for the first time that a neuropeptide expressed in a subpopulation of neurons is required for sleep homeostasis in a vertebrate. Our study supports a model in which neuronal activity is integrated to drive sleep pressure and in which Galanin-expressing neurons of the vertebrate hypothalamus play a critical role in coordinating sleep homeostasis.

## Results

### Short-term, pharmacologically induced increases in neuronal activity are followed by homeostatic rebound sleep in zebrafish

If the activity of neuronal assemblies across the brain is integrated to drive homeostatic rebound sleep, we hypothesized that short-term pharmacological increases of neuronal activity should accelerate homeostatic sleep pressure accumulation to trigger rebound sleep when the drugs are removed. The diurnal larval zebrafish is an ideal model to test this hypothesis because arousing drugs placed directly in the water are absorbed into the brain to rapidly modulate behavior, including sleep and wakefulness (Kokel et al., 2010; Rihel et al., 2010). Drug induced behavior is also rapidly reversible, as compounds can be quickly removed by washout (Mussulini et al., 2013; Tran et al., 2014). As a proof of concept, we first evaluated the effects of short-term exposure to the GABA-A receptor antagonist pentylenetetrazol (PTZ), which is known to potently and dose-dependently induce neuronal and behavioral hyperactivity in larval zebrafish (Baraban et al., 2005). Larvae raised and behaviorally tracked on a 14:10 light:dark cycle at 6 days post fertilization (dpf) were exposed to PTZ in the water for 1 hr in the morning, when homeostatic sleep pressure is low. Larvae treated with 10 mM PTZ increased their swimming activity and exhibited seizure-like behavior during the drug exposure, as has previously been described (Fig.1A, S1F). Following drug removal (washout), PTZ-treated larvae rapidly ceased to exhibit high frequency swim bouts (Fig. S1F) and entered a state of behavioral inactivity (rebound phase), which lasted until the end of the day/light period (Fig. 1A,B). Importantly, larval activity and sleep returned to normal control levels on the following day (Fig. S1 A,B).

**Fig. 1.**
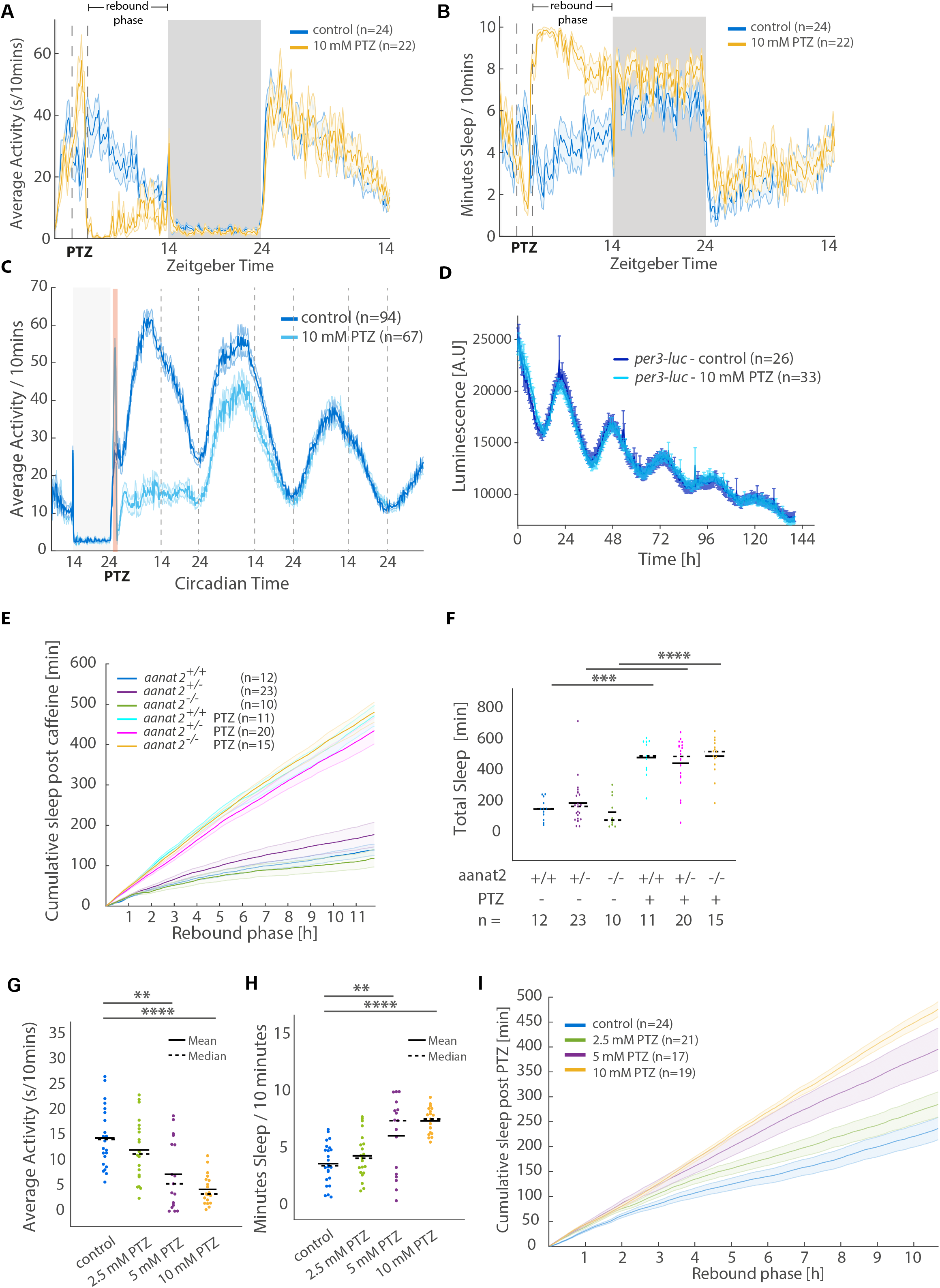
PTZ exposure is followed by rebound sleep. (A and B) A representative 36 hour tracking experiment showing that average activity is decreased (A) and average sleep is increased (B) after 1h of 10 mM PTZ exposure (rebound phase). Activity and sleep return to control levels the next day. The dashed line indicates timing of PTZ treatment, while the light and dark boxes indicate lights ON and OFF periods, respectively. The ribbon along each trace depicts ±SEM. The same representative experiment is quantified in G-I. Only the 10 mM PTZ behavior is plotted for clarity. (C) Activity behaviour trace following 1 hour PTZ exposure (red line) in constant light conditions. Dashed lines indicate subjective day/night transitions. Data is a combination of two independent experiments. (D) Luminescence measured as a result of rhythmic *per3* expression over circadian time post vehicle or PTZ treatment. (E) Cumulative rebound sleep during the rebound phase post 10 mM PTZ is not affected in *aanat2* mutant larvae. The ribbons of each cumulative trace depict ±SEM. (F) Quantification of sleep during rebound phase (E). Each dot represents a single larva. ***p<0.001, ****p<0.0001, Kruskal-Wallis, Dunn-Sidak post hoc (for all unless stated). (G and H) Quantification of activity and sleep during the rebound phase for control and three doses of PTZ indicate rebound sleep is dose-dependently affected. Each dot represents a single larva (see Fig. 1E for the n of each dose). **p<0.01, ***p<0.001, ****p<0.0001 (I) Cumulative rebound sleep is dose-dependently increased following exposure to increasing doses of PTZ. The ribbons of each cumulative trace depict ±SEM. See also Figure S1

To demonstrate that the inactive state observed following PTZ exposure is a *bona fide*, physiological sleep state, we tested whether post PTZ inactivity fulfils the key sleep behavioral criteria that have been previously established in zebrafish and other non-mammalian species (Hendricks et al., 2000; Prober et al., 2006; Raizen et al., 2008; Yokogawa et al., 2007; Zhdanova et al., 2001). Firstly, periods of continuous inactivity that last longer than one minute have been established as sleep in larval zebrafish and long rest bouts are rarely observed during the circadian light period when the diurnal zebrafish are predominantly awake (Prober et al., 2006; Rihel et al., 2010). Following exposure to PTZ, the total time spent asleep was significantly increased due to the lengthening of individual sleep bouts by an average of 1.7 fold and an increase in the number of sleep bout initiations (e.g. Fig. S2F). Secondly, sleep is accompanied by an increased arousal threshold to stimuli. Following 10mM PTZ-exposure, larvae showed lowered responsiveness to a series of mechano-acoustic tapping stimuli of varying strengths delivered during the rebound sleep phase (Fig. S1C) (Burgess and Granato, 2007; Singh et al., 2015). Third, unlike during paralysis or a coma, sleep is reversible with strong, salient stimuli. Following PTZ exposure, larvae responded to a series of 10 minute dark pulses with characteristic increases in locomotor behavior (Emran et al., 2007), similarly to controls (Fig. S1D), with the exception of the dark pulse immediately after drug washout. Furthermore, behavioral inactivity following PTZ-exposure was unlikely to be due to neuronal injury, as markers of apoptosis, including activated Caspase-3 (Fig. S1E, see also Fig. S5F) were not induced by drug exposure, and neither larval morphology nor sleep/wake behavior was altered one day following treatment (Fig. 1A, S1A,B). Together, these data indicate that inactivity following acute drug-induced neuronal hyperactivity is a sleep-like state comparable to physiological sleep.

To assess whether PTZ exposure affects circadian rhythmicity, we tracked larvae post PTZ over several days under constant light (LL) conditions, in which zebrafish larvae maintain molecular and behavioral rhythms without the drastic locomotor suppression of constant darkness. Although the suppression of activity following PTZ exposure was prolonged in LL compared to in LD, the drug had no effect on the circadian phase or period of waking behaviour (Fig. 1C). We also used a *per3-luciferase* transgenic zebrafish line (Kaneko and Cahill, 2005) to measure *per3* expression in constant dark following either vehicle or PTZ exposure and did not detect any changes to the amplitude, phase, or period of rhythmic *per3* expression (Fig. 1D). Moreover, the production of melatonin, which is required in zebrafish for the circadian rhythmicity of sleep (Gandhi et al., 2015), is not required for PTZ-induced rebound sleep as *aanat2* mutant larvae increase sleep after PTZ exposure to the same extent as both wild type and heterozygous siblings (Fig. 1E,F). We conclude that PTZ-induced rebound sleep is not due to alterations in circadian clock function.

Finally, to test whether PTZ-induced rebound sleep is under homeostatic control, we hypothesized that the amount of rebound sleep following PTZ exposure should correlate to the amount of PTZ administered, as the effects of PTZ on neuronal firing and behavioral hyperactivity have been shown to increase with increasing dose (Baraban et al., 2005; Ellis et al., 2012; Turrini et al., 2017). Indeed, the average activity and sleep of PTZ-treated larvae during the rebound phase varied dose-dependently (Fig. 1G,H), with total sleep accumulation increasing for many hours after drug exposure (Fig. 1I). Thus, in addition to fulfilling the key behavioral features of sleep, PTZ-induced rebound sleep is under homeostatic regulation.

### Pharmacologically induced rebound sleep is independent of physical hyperactivity, seizure induction, and pharmacological target

PTZ represents an extreme test case that induces high levels of physical and neuronal activity, including seizures. To assess that our observations were not the result of unwanted effects of PTZ, we tested additional, pharmacologically diverse drugs that promote neuronal activity and behavioral wakefulness in zebrafish larvae, including the adenosine receptor antagonist caffeine, the voltage-activated potassium channel antagonist 4-aminopyridine (4-AP), and the voltage-gated sodium channel agonist aconitine, one of the most potent modulators of behavioral wakefulness in zebrafish (Ellis et al., 2012; Rihel et al., 2010). Because these drugs have distinct pharmacological targets as well as diverse effects at the behavioral and neuronal levels, the features of drug-induced hyperactivity that are required to elicit rebound sleep could be dissected. Similar to PTZ, exposure to both caffeine and 4-AP for 1 hour induced significant increases in rebound sleep after drug removal (Fig. 2A,B; Fig. S2A,C,D-F) Following caffeine washout, cumulative sleep, rebound sleep bout length and number were dose-dependently increased (Fig. 2E-G). In contrast, aconitine failed to elicit rebound sleep despite inducing the most vigorous and extended behavioral activity during drug treatment at multiple doses (Fig. 1C,D,H-J, S2B-F). This is unlikely to be an effect of aconitine acting as an inhibitor of sleep homeostasis as treatment with a combination of aconitine and caffeine did elicit rebound sleep (Fig. S2G). Finally, we observed that only PTZ and 4-AP generated seizures, as measured by the number of high frequency swim bouts (Fig. S1F) or by calcium imaging (see below, Fig S3B). In contrast, caffeine failed to elicit seizures at the concentrations tested. Overall, these results indicate that pharmacologically induced rebound sleep is independent of physical activity, the presence of seizures, and pharmacological target, demonstrating that the degree of physical arousal during exposure does not predict rebound sleep behavior.

**Fig. 2.**
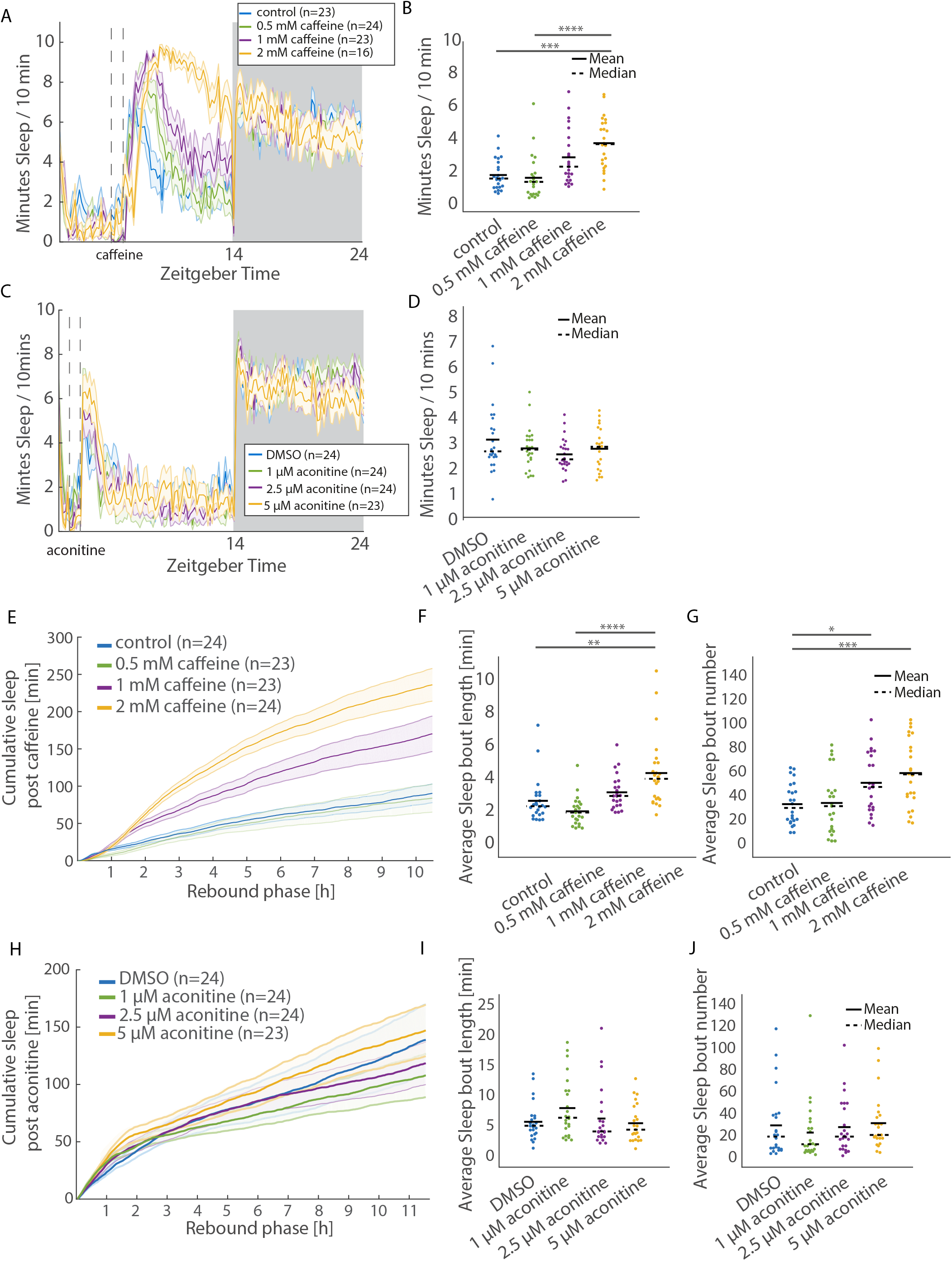
Caffeine but not aconitine exposure is followed by rebound sleep. (A and B) Sleep increased after 1h treatment with three doses of caffeine (rebound phase) (A). The dashed line indicates timing of caffeine treatment, while the light and dark boxes indicate lights ON and OFF periods, respectively. The ribbon along each trace depicts ±SEM. (B) Average data for individual larvae during the rebound phase is shown to the right of each behavior trace. **p≤0.01, ***p≤0.001, ****≤0.0001 (C and D) Sleep behavior remains unchanged after 1h treatment with three doses of aconitine (C). The dashed line indicates timing of aconitine treatment, while the light and dark boxes indicate lights on and off periods, respectively. The ribbon along each trace depicts ±SEM. Average data for individual larvae during the rebound phase is shown to the right (D). n.s. p>0.05 (E) Cumulative sleep after 1h exposure to caffeine (rebound phase) is dose-dependently increased. A single representative experiment is shown. The ribbons represent ±SEM. (F and G) Average sleep bout length and number are dose-dependently increased during the rebound phase following caffeine treatment. Each point represents a larva from the experiment in F. *p≤0.05, **p≤0.01, ***p≤0.001 ****p≤0.001 (H) Cumulative rebound sleep is unaffected relative to controls after 1h treatment with increasing doses of aconitine. The ribbons represent ±SEM. (I and J) Average sleep bout length and number are not significantly changed post aconitine during the rebound phase (Data from I and Fig S2C-D). See also Figure S2

### Pharmacologically induced rebound sleep is a function of global neuronal activity

Given that pharmacological target, presence or absence of behavioral seizures, and degree and duration of physical activity do not predict the ability of compounds to elicit post-washout rebound sleep, we speculated that the divergent response to these drugs could be explained by differential stimulation of neuronal activity. To examine this possibility, we measured the expression levels of the immediate early gene *c-fos*, a well-established correlate of neuronal activity (Baraban et al., 2005; Chung, 2015), directly after exposure to aconitine, caffeine, 4-AP and PTZ. All drugs that elicit rebound sleep— caffeine, 4-AP and PTZ— strongly induced *c-fos* mRNA expression in widespread neuronal populations (Fig. 3A,B). In contrast, aconitine, which fails to induce rebound sleep, increased *c-fos* only in a small region of the dorsal telencephalon, resulting in a low overall induction of *c-fos* expression (Fig. 3A,B). Importantly, aconitine did not affect the ability of caffeine to induce *c-fos* transcription (Fig. S3D). Total *c-fos* expression measured at the end of the drug exposure strongly correlated with the amount of subsequent rebound sleep that is induced across drugs (R^2^=0.96, linear regression) and as a function of both caffeine and PTZ dose (R^2^=0.99 and R^2^=0.93, linear regression, Fig. 3C-F). In contrast, neither the amplitude nor the duration of physical activity during drug exposure correlated with subsequent rebound sleep, suggesting that locomotion alone cannot account for homeostatic sleep drive (Fig. 3G-H; S3A). These results support the hypothesis that neuronal activity is integrated prior to homeostatic rebound sleep.

**Fig. 3.**
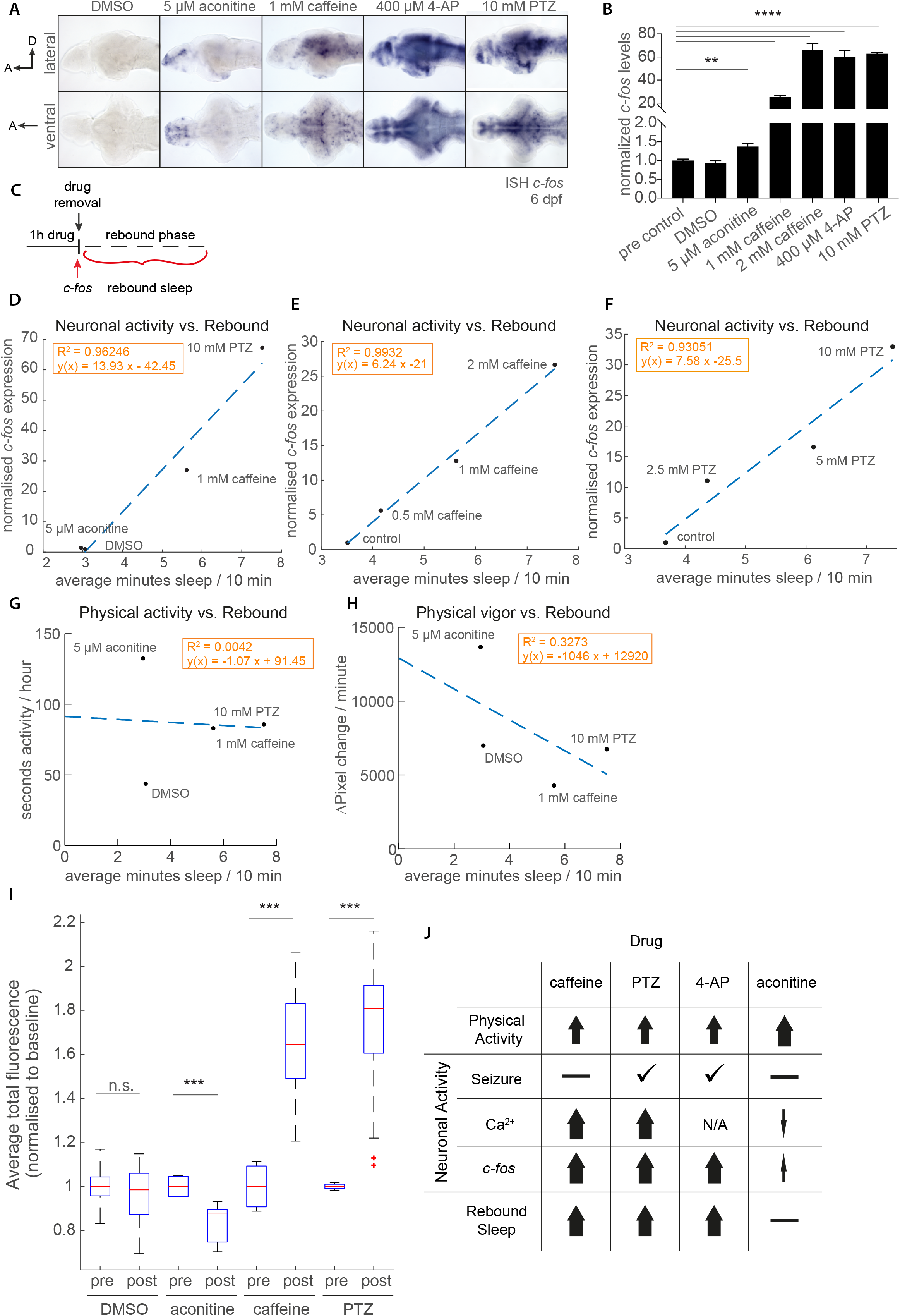
Global measures of neuronal, but not physical, activity correlate with subsequent rebound sleep amount. (A) ISH on dissected larval brains to visualise the immediate early gene, *c-fos* mRNA expression immediately after 1h exposure to aconitine, caffeine, 4-AP, and PTZ. Note that *c-fos* is broadly upregulated in many larval brain regions by rebound sleep-inducing compounds (caffeine, 4-AP, and PTZ) but is restricted to the telencephalon in aconitine treated larvae. A, anterior; D, dorsal. (B) *c-fos* expression measured by qRT-PCR over indicated drugs and doses. **p≤0.01, ****p≤0.001. All qRT-PCR measurements were performed on RNA extracted from 30-40 pooled larvae. (C) Schematic of experimental paradigm. Transcription levels of *c-fos* were determined by qRT-PCR on samples collected directly after 1h drug exposure. The rebound sleep was averaged over the rest of the circadian day (rebound phase). (D, E and F) The average sleep during the rebound phase is strongly correlated to the expression of *c-fos* induced by indicated drugs (D, R^2^= 0.96), increasing doses of caffeine (E, R^2^=0.99) and increasing doses of PTZ (F, R^2^=0.93). The dashed line depicts the linear regression curve. (G and H) There is no correlation between average rebound sleep and the duration of movement (G, R^2^=0.004) and a slight negative correlation with the magnitude of movement (H, R^2^=0.33) in response to the indicated drugs. The dashed lines depict the linear regression curve. (I) Whole brain fluorescent Ca^2+^ imaging before and after addition of pharmacological compounds shows that only PTZ and caffeine induces increases in neuronal activity signals over baseline, as measured by changes in average fluorescence. Averages of two (pre) or seven (post) 5min recordings of three larvae are shown. Brains were imaged at 5.5 fps and signals normalised to the average baseline recorded pre drug. Boxes indicate 25^th^ and 75^th^ percentiles, and whiskers extend to most extreme data points not considered to be outliers (i.e. 1.5 times the interquartile range away). ***p≤0.001. Wilcoxon signed rank test. (J) Summary of wake-promoting drugs’ effects on physical activity, seizure induction, and neuronal activity as measured by *c-fos* expression and whole-brain calcium imaging. Only measures of neuronal activity are consistent with the induction of post-exposure rebound sleep.

While immediate early genes such as *c-fos* are widely used as correlates of neuronal activity, some active neurons do not induce *c-fos*, and other neurons exhibit inconsistent *c-fos* transcriptional induction during repeated rounds of enhanced activity (Kovacs, 2008). To corroborate the *c-fos* measurements of drug-induced neuronal activity, we turned to *in vivo* wide-field Ca^2+^ imaging as a second, more direct readout of neuronal activity (Turrini et al., 2017). Using Tg(elav3:H2B-GCaMP6s) transgenic larvae, which express the pan-neuronal, nuclear-localized genetically-encoded Ca^2+^ indicator GCaMP6s (Freeman et al., 2014), we measured total fluorescence changes across the whole brain over a 1 hour exposure to DMSO, 5 µM aconitine, 2 mM caffeine, and 10 mM PTZ (see Extended Methods). In agreement with the *c-fos* expression measurements, PTZ and caffeine, but not the DMSO control, caused a measureable increase in total fluorescence above baseline, while aconitine caused a global decrease in activity (Fig. 3I; S3B-C, Supplemental Movies 1-3). Additionally, although 2mM caffeine upregulated neuronal activity in many areas of the brain, only PTZ produced detectable seizures, observed as large, rapid, and synchronized increases in fluorescence followed by a rapid decrease in global fluorescence levels (Fig. S3B-C and Supplemental Movies 1-3).

Together, these data indicate that increases in global measures of neuronal activity during drug-induced wakefulness vary across drugs and predict the amount of subsequent rebound sleep after drug removal, whereas neither the presence of seizures nor the intensity of physical activity correlate with rebound sleep amount (Figure 3J).

### Galanin expressing neurons of the preoptic area are active during rebound sleep

Knowing the identity of candidate sleep-control neurons that are active during rebound sleep might provide clues to how brain-wide neuronal activity during waking is integrated to generate the appropriate amount of subsequent rebound sleep. We therefore turned to whole-brain activity mapping to identify rebound sleep-active neurons following pharmacologically induced neuronal activity. We used mitogen activated protein kinase (MAP) – mapping (Randlett et al., 2015), which has been shown to be a reliable readout of neuronal activity in zebrafish larvae, to create whole brain neural-activity maps 4 hours into pharmacologically induced rebound sleep. Larval brain activity maps during post-PTZ (Fig. 4A) or post-caffeine (Fig. 4C) rebound sleep showed several areas with altered activity (FDR threshold: <5×10^−5^) compared to control brains, which were identified by morphing onto the Z-brain reference atlas (Supplemental Table 1). Areas with a significant increase in pERK/tERK activity during rebound sleep induced by both drugs include the neuropil and stratum periventriculare of the optic tectum, the cerebellum (both the lobus caudalis and corpus cerebelli), the torus semicircularis, the intermediate hypothalamus, the myelencephalic choroid plexus and the pineal gland (Fig. 4A,C). A few areas also showed relative downregulation of pERK/tERK during rebound sleep, including the olfactory bulb dopaminergic cluster and the subpallial Gad1b cluster (Supplemental Table 1).

**Fig. 4.**
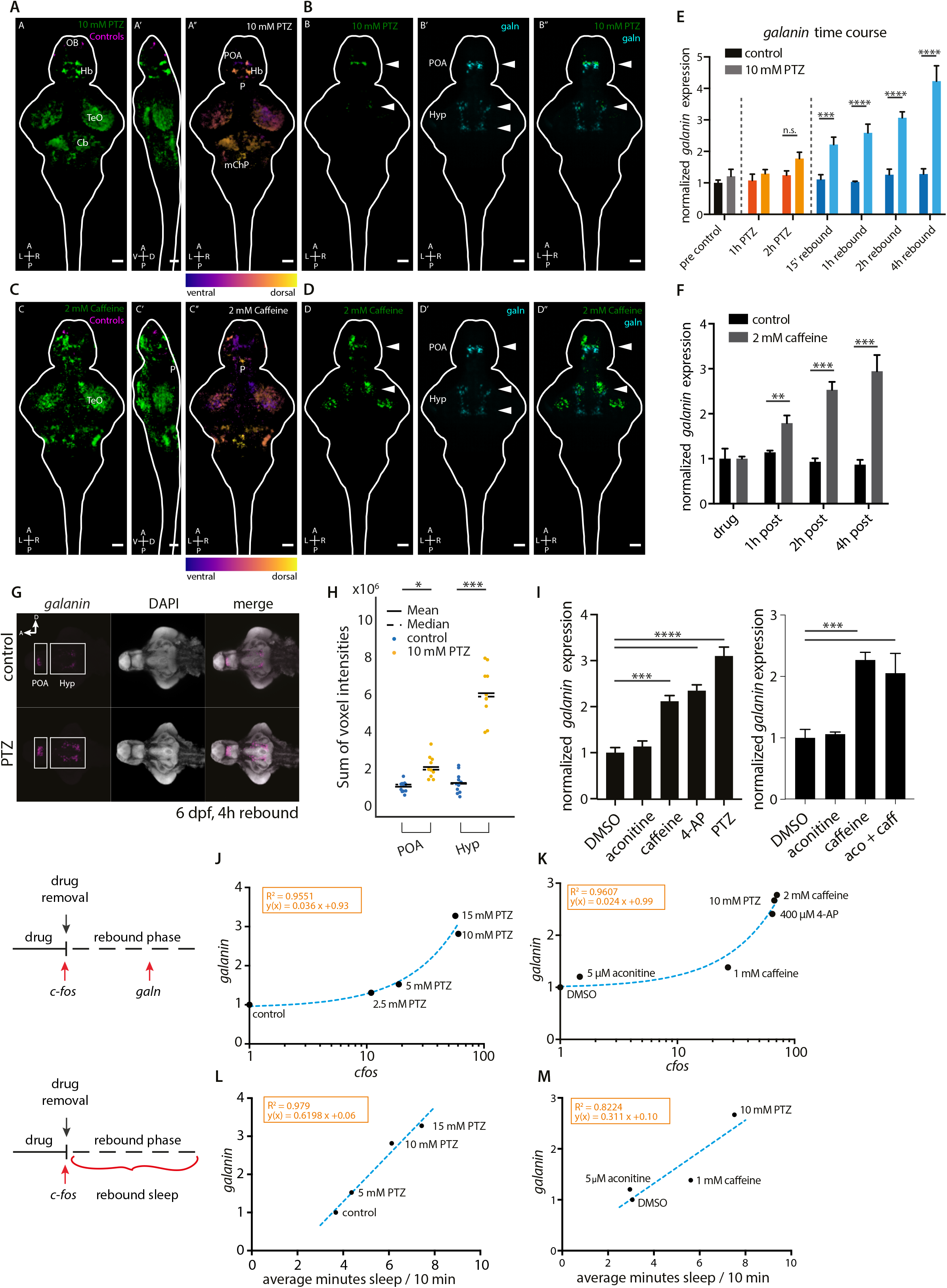
Whole brain neural-activity mapping and *galn* transcriptional analysis. (A,A’) Dorsal and lateral MAP-maps obtained from 6 dpf larvae exposed to 10 mM PTZ (n=24) or control (n=24) for 1h and allowed to recover for 4h after washout. Whole brain maximum-intensity projections of the MAP-map from 10 mM PTZ versus controls. Green and magenta indicates areas up- and down-regulated during PTZ-induced rebound sleep versus controls, respectively. (A’’) Depth projection of the positive MAP-map from 10 mM PTZ versus controls (blue=ventral, yellow=dorsal). (B-B’’) A 60 µm-thick maximum-intensity projections of the ventral larval brain from the MAP-map of 10 mM PTZ versus controls (B), of a *galn* fluorescent ISH in a 6 dpf brain morphed to the same reference brain (B’), and of the merged signals (B’’), showing the overlap between the upregulated region in the POA and hypothalamus and the *galn* neurons in these areas (Arrowheads). (C,C’) Dorsal and lateral MAP-maps obtained from 6 dpf larvae exposed to 2 mM caffeine (n=22) or control (n=24) for 1h and allowed to recover for 4h after washout. Whole brain maximum-intensity projections of the MAP-map showing brain areas relatively upregulated during rebound sleep after exposure to 2 mM caffeine (green) or in controls (magenta). (C’’) Depth projection of the positive MAP-map from 2 mM caffeine versus controls (blue=ventral, yellow=dorsal). D-D’’) A 60 µm-thick maximum-intensity projections of the larval brain from 2 mM caffeine versus controls (D), of a *galn* fluorescent ISH (D’), and of the merged signals (D’’), showing a similar overlap between the upregulated region in the POA and the POA *galn* neurons (Arrowheads). Cb, cerebellum. mChP, myelencephalic choroid plexus. Hb, habenula. Hyp, hypothalamus. OB, olfactory bulb. P, pineal. POA, preoptic area. TeO, tectum opticum. A, anterior; P, posterior; R, right; L, left; D, dorsal; V, ventral. Scale bars, 50 µm (all panels). (E) *galn* qRT-PCR time course including pre-exposure control samples (black/grey), 1h and 2h PTZ exposure (red/orange) and indicated time points during the rebound sleep phase (dark/light blue). Controls depicted in the darker shades are time-matched to PTZ-treated samples. n.s. p>0.05, ****p≤0.0001. One-way ANOVA, Bonferroni corrected for multiple comparisons. (F) *galn* qRT-PCR time course in response to 1h, 2 mM caffeine. Timed controls (black) compared to treated samples collected at the indicated time points during the rebound sleep phase (grey). ***p≤0.001, ****p≤0.0001. One-way ANOVA, Bonferroni corrected for multiple comparisons. (G) Fluorescent ISH comparing *galn* expression (magenta) in control and PTZ treated larval brains 4h into the rebound sleep phase. Nuclei are labelled with DAPI (grey). White boxes outline the region of interest (ROI) for the preoptic area (POA) and hypothalamic (Hyp) neuron populations, which were used to calculate intensity in H. Scale bar: 100 microns (H) Intensity of fluorescent signal is compared between the control and PTZ treated larvae during rebound phase for the POA and Hyp *galn* populations (H). Each dot is a measurement for individual larvae (n=12). *p≤0.05, ***p≤0.001. (I) Bar graph showing qRT-PCR quantification of mean *galn* mRNA expression after treatment with indicated pharmacological compounds normalized to vehicle control 2.5h into the rebound sleep phase. Error bars indicate the standard deviation (n=3). ***p ≤ 0.001, ***p ≤ 0.001, ****p ≤ 0.0001, one-way ANOVA with post hoc Dunnett’s test for comparisons to DMSO-treated controls. (J and K) Correlation between *c-fos* expression (x-axis, note log scale) measured directly after exposure to increasing concentrations of PTZ (J) or indicated pharmacological compounds (K) and *galn* expression 4h into rebound phase (y-axis). The dashed lines depict linear regression curves (R^2^=0.96 in J, R^2^=0.96 in K). (L and M) Correlation between average rebound (x-axis) sleep post PTZ (L) or indicated pharmacological compounds (M) and *galn* expression 4h into rebound phase (y-axis). The dashed lines depict linear regression curves (R^2^=0.98 in L, R^2^=0.82 in M). See also Figure S3 and Table S1

One set of neurons that were activated during both PTZ and caffeine induced rebound sleep were found in the preoptic area (POA) (Fig. 4A,C, Supplemental Table 1), the zebrafish homolog of the mammalian ventro-lateral preoptic area (VLPO), which contains sleep-active neurons that express the inhibitory neuropeptide Galanin (Galn) (Kroeger et al., 2018). We morphed a *galn* fluorescent ISH image onto the same reference brain and found a strong overlap in sleep-active pERK/tERK signals and *galn* expression in the POA (Fig. 4B,D). In addition, although the resolution of MAP-mapping for sparsely, non-stereotypically distributed neurons is more limited, we observed increased neuronal activity in the anterior hypothalamus, where *galn* is also expressed in zebrafish (Fig. 4B,D). These data indicate that the homolog of at least one important sleep-active centre of the mammalian brain, the Galn expressing VLPO, is also active during rebound sleep in zebrafish.

### *galn* expression correlates with rebound sleep pressure

Increased neuronal activity is often accompanied by changes in neurotransmitter transcription rates via ‘stimulus-secretion-synthesis coupling’ (MacArthur and Eiden, 1996), presumably as a mechanism to replenish neuropeptide stores after secretion. We therefore tested whether the increased activity of Galn neurons during rebound sleep was accompanied by upregulation in *galn* transcription. Indeed, *galn* expression increased steadily to 3-4 fold after both PTZ and caffeine washout during the rebound sleep phase (Fig. 4E,F), and returned to baseline the next day (Fig. S3I), consistent with a return of sleep behavior to control levels (Fig. S1A,B). Quantification of *galn* signal intensity using fluorescent *in situ* hybridization (ISH) revealed that both the preoptic and hypothalamic sub-populations selectively upregulate *galn* expression (Fig. 4G,H). The upregulation of *galn* transcription was limited to drugs that elicit rebound sleep (PTZ, caffeine, and 4-AP), as both qRT-PCR and ISH found no induction of *galn* after DMSO or aconitine exposure, and aconitine does not affect the ability of caffeine to induce *galn* (Fig. 4I, S3H). Moreover, *galn* expression is not under circadian regulation, as expression in constant light conditions is constant over circadian time, unlike the canonical circadian gene *per3* (Fig. S3E).

If *galn* expression is increased during rebound sleep as a function of heightened sleep pressure generated by increased neuronal activation, we predicted that the magnitude of *galn* induction would reflect both the magnitude of whole-brain neuronal activity generated during drug exposure as well as the total amount of subsequent rebound sleep. Indeed, *galn* transcription during rebound sleep is highly correlated with the dose-dependent increase in *c-fos* after PTZ exposure (R^2^=0.96, linear regression) as well as across drugs (R^2^=0.96, linear regression) (Fig. 4J,K). Moreover, *galn* expression levels are a good readout of the amount of rebound sleep induced by both PTZ and across drugs (R^2^=0.98 and 0.82 respectively, Fig. 4L,M). In contrast, *galn* transcription is uncorrelated to both the magnitude and the duration of physical activity (Fig. S3F,G), further supporting our evidence that neuronal but not physical activity contributes to homeostatic rebound sleep.

### *galn* mutants have reduced sleep but no increased seizure susceptibility

Whole brain activity mapping and *galn* expression analysis suggest that Galn-expressing neurons are recruited during pharmacologically induced rebound sleep in response to increased sleep pressure. To further examine the role of *galn* in regulating rebound sleep states, we used Crispr-Cas9 to generate two zebrafish *galn* mutant alleles targeting the 3^rd^ exon (Fig. S4A,B; see Extended Methods). Both are likely functionally null alleles of *galn*, as *galn* transcripts are greatly reduced, an indicator of nonsense-mediated decay (*galn*^*int3*^; Fig. S4C), and Galn peptide is not detectable by whole mount immunohistochemistry (*galn*^*i8*^; Fig. S4D). Both alleles are homozygous viable and fertile as adults, and they lack overt morphological phenotypes, with the exception of a loss of yellow-pigmented xanthophores (Fig. S5A,B).

Both homozygous *galn*^*i8/i8*^ and *galn*^*int3/int3*^ larvae show small but consistent disruptions in their sleep/wake structure compared to their wild type and heterozygous siblings when tracked in a 14:10 light:dark cycle over several days (Fig. 5A,B, and Fig. S4E-G). During the day, both *galn ^int3/int3^* and *galn*^*i8/i8*^ mutants have a significant reduction in sleep compared to sibling controls, due to both the initiation of fewer and shorter sleep bouts (Fig. 5B, S4E). Daytime sleep latency (i.e. time until the first rest bout from lights ON) is also increased (Fig. 5B, S4E) and average activity as well as waking activity during the day is higher in both *galn*^*int3/int3*^ and *galn*^*i8/i8*^ mutants. Total sleep and average sleep bout lengths at night are significantly but modestly reduced in *galn*^*int3/int3*^ (Fig. 5B, S4E), suggesting that *galn* is largely dispensable or redundant for maintaining sleep during the circadian night. Moreover, circadian rhythmicity is not disrupted in *galn* mutants, as we did not detect changes in the amplitude, phase, or periodicity of *per3* expression or sleep/wake behaviour compared to wild type siblings (Fig. S5C,D).

**Fig. 5.**
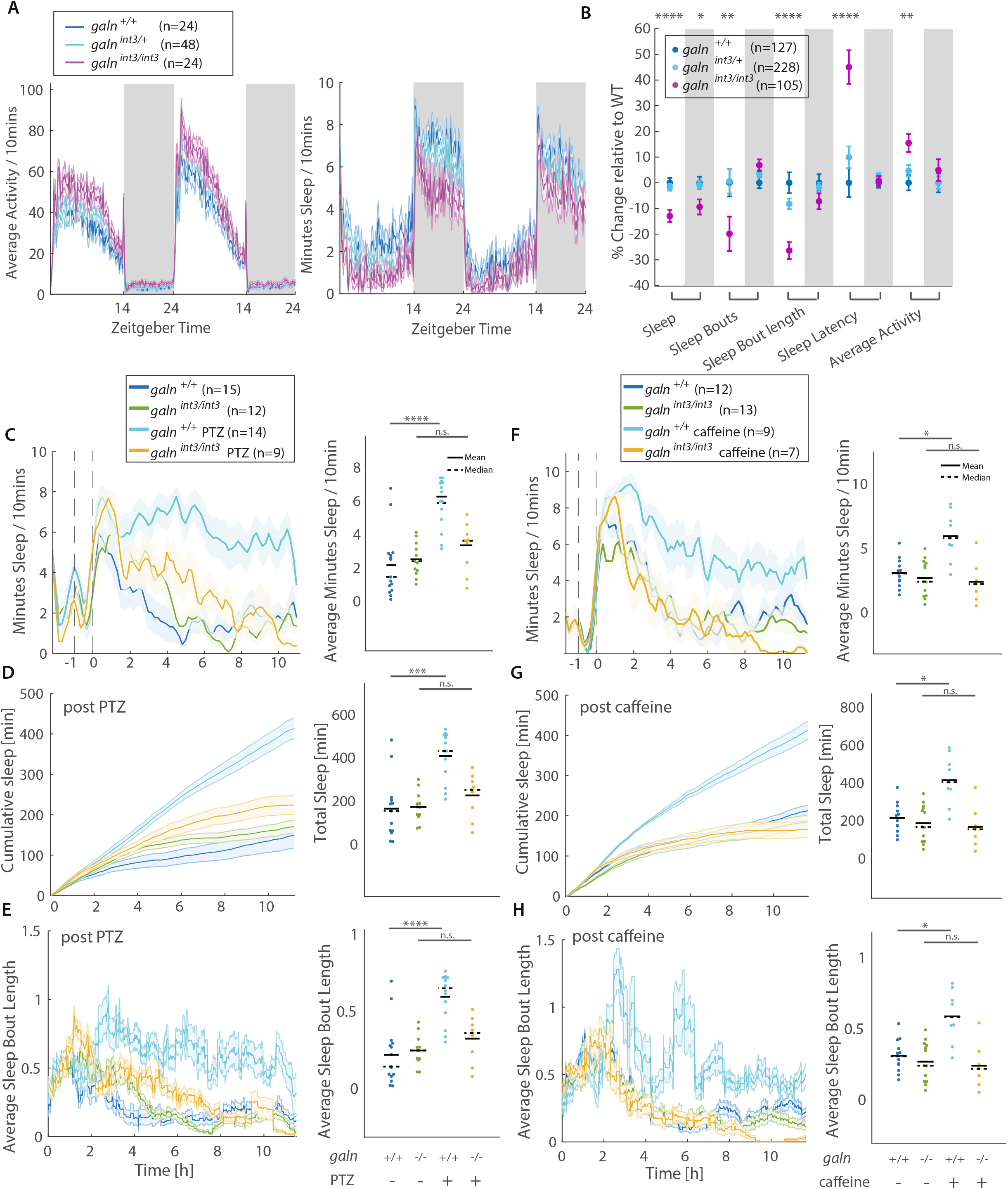
Galn is required for neural activity-induced rebound sleep. (A) Locomotor and sleep behavior traces for a representative experiment. White and grey areas indicate lights-ON and lights-OFF periods, respectively. The ribbons depict ±SEM. (B) Several features of larval sleep architecture during the day (white background) and night (grey background) are plotted for *galn*^*int3/int3*^ mutants as the mean % change relative to their wild type siblings. Error bars indicate ±SEM from four independent, blinded experiments. *p≤ 0.05, **p<0.01, ***p≤ 0.001, ****p≤ 0.0001 (C-H) Behavioral analysis of *galn*^*int3/int3*^ mutants and their wild type siblings following 5 mM PTZ (C-E) and 1 mM caffeine (F-H) exposure. For clarity *galn*^*int3i+*^ larvae are not shown. (C,F) depict the 30 min running average of sleep amount to the left and mean rebound sleep for each larva to the right. The start (t= −1) and end (t=0) of drug exposure are indicated with dashed lines. Neither PTZ nor caffeine treated *galn*^*int3/int3*^ mutants show significant changes in sleep (C and F) during the rebound phase (t≥0) relative to un-exposed *galn*^*int3/int3*^ mutants and their wild type siblings. (D and G) Cumulative sleep during the rebound phase following drug exposure is plotted as the mean ± SEM. Scatter plots quantify the total rebound sleep for each larva. (E and H) The average sleep bout length (ribbons depict ±SEM) during the rebound phase is plotted as a 60 min running average. Only wild type and not *galn*^*int3/int3*^ larvae lengthen sleep bouts during the rebound phase following exposure to either PTZ or caffeine. *p≤ 0.05 ***p≤ 0.001, ****p≤ 0.0001, n.s. p>0.05 See also Figure S4 and 5

Galn has been reported to play a neuroprotective role, for example after seizure and neuronal injury (Elliott-Hunt et al., 2004). To test if *galn* mutants exhibit increased neuronal apoptosis after seizures, we stained *galn*^*int3/int3*^ mutants and *galn*^*+/+*^ larval brains for activated Caspase-3 after exposure to 10 mM PTZ. Neither *galn*^*+/+*^ nor *galn*^*int3/int3*^ larval brains showed increases in activated Caspase-3 signal at the PTZ dose applied (Fig. S5F). Galn has also been ascribed antiepileptic properties (Podlasz et al., 2018). To assess seizure susceptibility in *galn*^*int3/int3*^, we treated larvae with an intermediate low dose of PTZ and counted high amplitude swim bouts as a measure of behavioral seizures (Baraban et al., 2005). *galn*^*int3/int3*^ mutants did not exhibit spontaneous behavioral seizures at baseline and responded similarly as *galn*^*+/+*^ larvae to PTZ (Fig. S5G), indicating that, unlike other seizure-susceptible zebrafish mutants (e.g. *cntnap2*; (Hoffman et al., 2016)), *galn* mutants are not more sensitive to the convulsant effect of PTZ.

### Galn is required for homeostatic rebound sleep after both pharmacologically induced neuronal activity and sleep deprivation

To test whether *galn* is necessary for pharmacologically induced rebound sleep, *galn*^*int3/int3*^, *galn*^*int3/+*^ and *galn*^*+/+*^ siblings were exposed to 1 hour of vehicle, PTZ or caffeine and behaviorally tracked for rebound sleep after washout. As expected, *galn*^*+/+*^ exhibited prolonged rebound sleep following PTZ, leading to a net accumulation of 407 min of sleep vs. 163 min in controls due to longer sleep bout lengths (Fig. 5C-E). In contrast, *galn*^*int3/int3*^ mutants had no statistically significant (p>0.05) increase in rebound sleep compared to controls (224 vs. 171 min) (Fig. 5C-E, S4H). Similarly, caffeine induced no rebound sleep in *galn*^*int3/int3*^ relative to vehicle control larvae (165 vs. 185 min), while *galn*^*+/+*^ siblings had significant increases in sleep following caffeine exposure (413 vs. 213 min) (Fig.5F-H, Fig. S4I). Induction of *c-fos* in response to PTZ and caffeine is not altered in *galn* mutants (Fig. S5E). Together, these data indicate that *galn* is required for rebound sleep after the pharmacological induction of neuronal activity.

Although pharmacologically induced neuronal activity triggers rebound sleep in larval zebrafish that satisfies the behavioral criteria of sleep, the levels of neuronal activity acutely induced by our assay are much higher than what animals naturally encounter acutely. To test whether pharmacologically induced, Galn-dependent rebound sleep is a general mechanism relevant to sleep homeostasis, we sleep deprived *galn* mutants by physically extending wakefulness throughout the night. To do this, we developed an optomotor response (OMR) “treadmill” sleep deprivation (SD) assay, in which larvae are sleep deprived by presenting moving black/white stripes, a strongly salient position stabilizing reflex that induces persistent swimming (Fig. 6A, see Extended Methods). The control group was exposed to a stationary version of the same grating to account for environmental effects, such as changes in luminance. Exposure to OMR during the day had no effect on wild type sleep compared to freely exploring larvae (Fig. 6B), suggesting that physical exhaustion is not a confounding factor in this assay. In addition, *c-fos* expression was actually reduced during OMR compared to freely exploring larvae during the day (Fig. S6A,B), which is consistent with the inability of daytime OMR to induce rebound sleep.

**Fig. 6.**
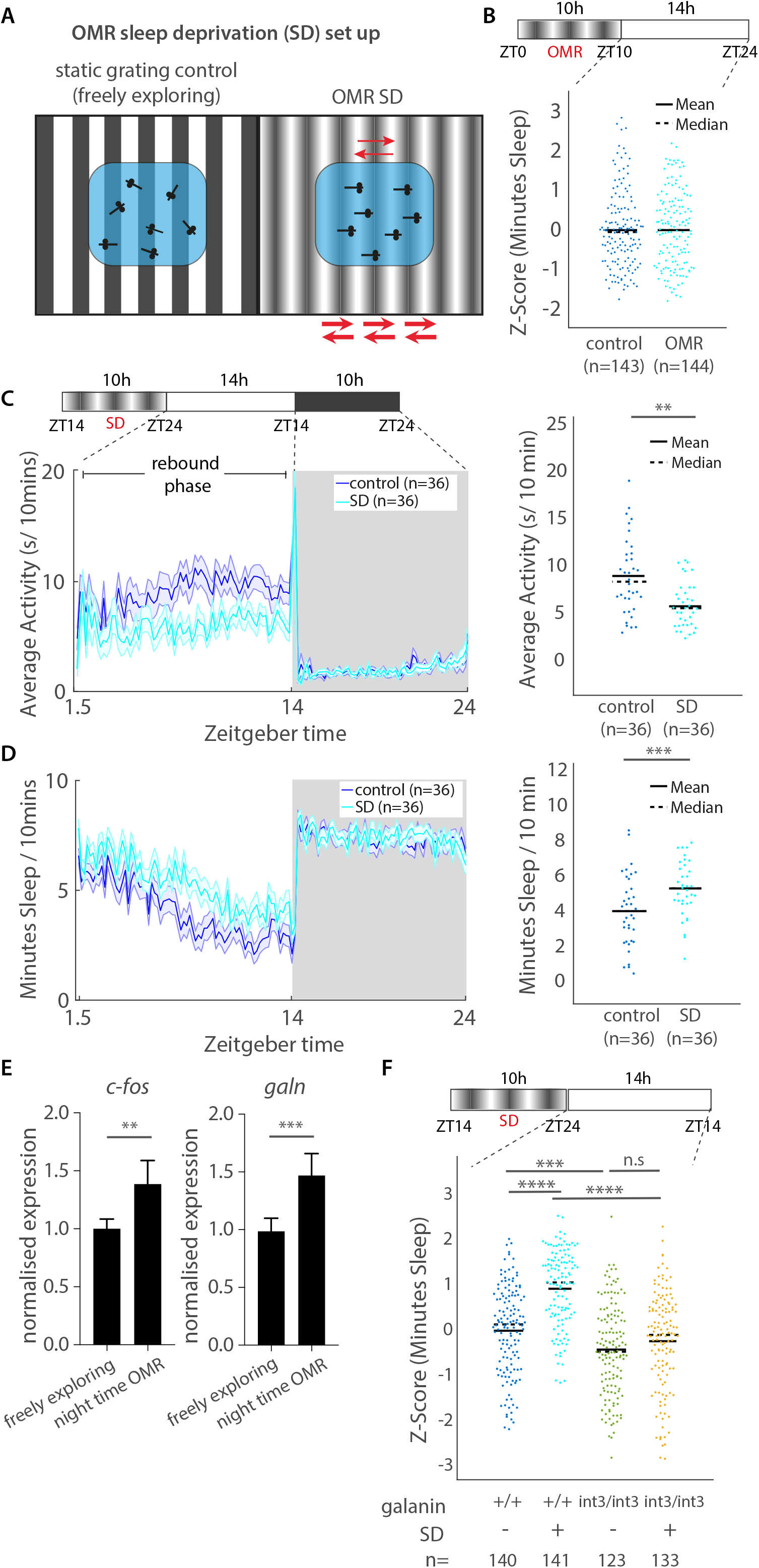
Galn is required for rebound sleep after sleep deprivation. (A) OMR sleep deprivation (OMR SD) setup consists of a moving and non-moving stripe condition. (B) Exposure to the OMR stimulus for 10h during the circadian day (ZT0-ZT10) does not affect subsequent sleep behavior. Each dot represents a single larva normalized to the mean control sleep amount (Z-score) across 3 independent experiments. ns=p>0.05 (C and D) Locomotor activity and sleep traces following a representative OMR SD experiment. The average activity and sleep are quantified to the right. Dots represent individual larva **p≤0.01, ***p≤0.001 (E) Bar graphs showing the mean ±SEM of RT-qPCR measured *c-fos* and *galn* expression after OMR sleep deprivation. **p<0.01, ***p≤0.001. one-way ANOVA. n = independent biological replicates. (F) Behavioral analysis of *galn*^*int3/int3*^ mutants following OMR SD shows that the increase in sleep following OMR SD of wild type siblings does not occur in *galn*^*int3/int3*^ larvae. Each dot represents a single larva normalized to the mean control sleep amount (Z-score) and combines 3 independent experiments. Two-way ANOVA (genotype x condition, p=2.8×10^−05^) followed by Tukey-Kramer post-hoc test. n.s. p>0.05, ***p≤ 0.001, ****p≤ 0.0001. See also Figure S6

In contrast, OMR SD over a 10 hour deprivation period spanning the night, when larval sleep is most prevalent, was followed by increased sleep on the subsequent day compared to un-deprived controls (Fig. 6C,D). In agreement with data from our pharmacological assay, *c-fos* expression during the night-time OMR SD was increased relative to sleeping controls and *galn* expression was subsequently upregulated during the rebound sleep phase (Fig. 6E). *galn*^*+/+*^ but not *galn*^*int3/int3*^ mutants showed significant rebound sleep after one night of sleep deprivation (Fig. 6F), suggesting that *galn* is required for the increased rebound sleep following extended wakefulness by OMR SD. Importantly, this was not due to *galn*^*int3/int3*^ mutants losing less sleep than siblings during OMR SD(Fig. S6C). Taken together, these results show that rebound sleep following both pharmacologically induced neuronal activity and sleep deprivation engages *galn*, which is required for homeostatic rebound sleep in both paradigms.

## Discussion

### Decoupling sleep pressure from prior waking implicates global neuronal activity as a driver of sleep homeostasis

Measures of sleep pressure, including the magnitude of cortical SWA as well as the duration of rebound sleep, increase as a function of prior waking time (Borbely and Achermann, 1999; Borbely et al., 1981; Dijk et al., 1987; Dijk et al., 1990). Most studies challenge the sleep homeostat through protocols that enforce extended wake. Our pharmacological sleep homeostasis assay, which converges onto the same regulatory mechanisms as physical sleep deprivation, allowed us to decouple sleep pressure from prior waking duration. Instead of waking time, physical vigor, or total active duration, we found that global measures of neuronal activity were a potent predictor of subsequent homeostatic rebound sleep. We therefore propose that, in addition to the prior time spent awake, sleep pressure is strongly influenced by prior waking intensity as determined by the recruitment of neurons during wakefulness.

Although rarely tested directly, the concept that the quality of waking activity contributes to sleep pressure is consistent with some previous experimental data in other species. In rats, exposure to a 1 hour social stress increased subsequent SWA to a similar degree as a 12 hour sleep deprivation (Meerlo et al., 1997). Sleep deprivation techniques in mice that induce higher levels of *c-fos* expression, such as continual cage changes versus gentle handling, also build more sleep pressure as reflected in SWA during rebound sleep (Suzuki et al., 2013). In humans, exposure to stimulating environments decreased sleep latency and increased sleep depth relative to stereotypical conditions (Horne and Minard, 1985). More extremely, EEG recordings from patients suffering from chronic seizures revealed higher levels of SWA compared to healthy controls (Kaltenhauser et al., 2007; Schonherr et al., 2017), and post-ictal fatigue is a common symptom in epilepsy patients (Ettinger et al., 1999; Hamelin et al., 2010).

Decoupling physical activity and total waking time from the sleep homeostasis machinery may also provide a mechanistic basis by which animals avoid rebound sleep pressure under extreme environmental or behavioral fitness pressures. For example, starved or courting flies significantly increase foraging or mating behavior with little apparent rebound sleep (Beckwith et al., 2017; Keene et al., 2010), sandpiper males during mating season exhibit insomnia lasting days with little build-up of sleep pressure (Lesku et al., 2012), and killer whale newborns and their mothers show almost no sleep during the first weeks after birth (Lyamin et al., 2005). Perhaps under exceptional conditions, a kind of “behavioral autopilot” is engaged to minimize the recruitment of neurons to high levels of activity and thus prevent the accumulation of sleep pressure. Consistent with this concept, we found that OMR-induced swimming during the wake period induced less neuronal activity than freely exploring larvae. Similarly, mice engaged in a stereotypical running wheel behavior show reduced firing rates in motor- and somatosensory cortical neurons and are capable of longer wake times (Fisher et al., 2016).

Our results favour a model in which the intensity of global neuronal activity is integrated to drive sleep pressure, with brain-wide measurements of neuronal activity accurately predicting the duration of subsequent rebound sleep. This relationship of neuronal activity to sleep homeostasis holds both across doses of drugs with different mechanisms of action and during physical sleep deprivation. However, consistent with our data is the idea that only a few critical neurons— perhaps even Galn-expressing neurons themselves—are required to be active during wakefulness to trigger rebound sleep. This “privileged” neuron hypothesis has been suggested in *Drosophila*, in which the thermogenic activation of distinct wake-promoting neurons that drive equal amounts of sleep loss induce various levels of subsequent rebound sleep (Seidner et al., 2015). Whether these results reflect the total number of neurons recruited during waking or the activation of a critical few neurons will require exploration with simultaneous behavioral tracking, whole-brain neuronal imaging, and optogenetic and chemogenetic manipulation of circuits both during and after sleep deprivation.

### Galn as an output arm controlling homeostatic rebound sleep

Our transcriptional, neuronal activity, and genetic lesioning data in zebrafish converge on *galn* as sensitive to increases in sleep pressure and Galn being required for the induction of subsequent rebound sleep. These results not only demonstrate a critical role for the neuropeptide Galn in regulating sleep homeostasis but also identifies a key neuronal subpopulation upon which sleep pressure must ultimately act to drive rebound sleep.

In mammals, the Galn-expressing, GABAergic neurons of the ventrolateral preoptic area (VLPO) have long been proposed to be a major sleep regulator, as up to 80% of sleep active neurons in the VLPO are Galn positive (Gaus et al., 2002) and lesioning the VLPO leads to insomnia in the rat (Lu et al., 2000). However, the specific role of Galn neurotransmission in sleep has not been established (Saper and Fuller, 2017), with most studies having focused on the function of galaninergic neurons rather than the neuropeptide itself. For example, chemo- and optogenetic stimulation of VLPO Galn-expressing neurons increases sleep (Kroeger et al., 2018) (although see (Chung et al., 2017)), but these experiments cannot distinguish among the roles of GABA, Galn, or other transmitters. To our knowledge, the role for Galn in sleep homeostasis has not been tested before in mammals, probably because mouse *Galn* knockouts have pleiotropic phenotypes, including the inability to lactate (Wynick et al., 1998), pain sensitivity (Kerr et al., 2000), and alterations in feeding (Adams et al., 2008), consistent with the distribution of Galn expression in multiple neuronal subpopulations. In contrast, the expression of *galn* is restricted to fewer neurons in larval zebrafish, and the sleep behavioral assay is performed prior to feeding and other complex behaviors, which perhaps allowed us to uncover *galn*’s conserved role in sleep. However, unpublished data showing that ablation of the preoptic Galn-expressing neurons in mice leads to the loss of rebound sleep (Ma et al., bioRxiv) indicates that the role of Galn in regulating sleep homeostasis is not likely limited to zebrafish and is conserved across vertebrates.

Using two separate assays, we demonstrated that *galn* is required for converting sleep pressure into changes in homeostatic sleep behavior. In flies, the sleep promoting neurons that project to the dorsal fan-shaped body of the central complex, which has been likened to the mammalian VLPO, use the insect specific neuropeptide Allatostatin for inhibition of arousal and have been suggested to act as a gate to regulate homeostatic sleep need (Donlea et al., 2018). A similar mechanism could apply in zebrafish, in which sleep pressure increases as a result of neuronal activity to dose-dependently regulate the activity or excitability of *galn* neurons and/or the release of Galn. Consistent with this idea, in rodents, the sleep-active Galn neurons of the VLPO increase their firing rate in response to sleep deprivation and during rebound sleep, with firing returning to baseline levels as SWA decreases (Alam et al., 2014). Future experiments will be required to tease out the specific roles of fast GABAergic neurotransmission versus slower release of the neuromodulator Galn in this process, but one possibility is that rapid neurotransmitter release signals sleep initiation, while Galn release is required to stabilize or maintain recovery sleep after deprivation.

Our study significantly expands the current understanding of homeostatic sleep regulation by demonstrating that *galn* is required to generate appropriate rebound sleep. Important next steps include the identification of the molecular and cellular signals upstream of Galn that link neuronal activity to sleep homeostasis and elucidating the downstream circuits that are directly modulated by *galn* neurons to orchestrate rebound sleep behavior. Another question for future studies is to uncover potential interactions between *galn* and other putative sleep homeostasis genes, for example using Galn-expressing and other neuron specific genetic knockouts to determine if these circuits are directly or indirectly impacted by the loss of sleep homeostasis genes. Finally, it is likely that Galn is not the sole regulator of homeostatic sleep, as *galn* mutants retain some homeostatic responses after very strong arousing stimuli (e.g. after 2mM caffeine exposure, Fig. S4I). One potential regulator that should be examined in this context is the sleep-inducing neuropeptide Prokineticin2, which can upregulate *galn* transcription in zebrafish (Chen et al., 2017). The modest reduction of normal sleep at night in *galn* mutants under baseline conditions also points to other, as yet undiscovered, sleep regulators and hints that baseline and homeostatically-driven rebound sleep may have distinct modifying pathways. Although many questions remain, the simplicity and scalability of our pharmacologically induced rebound sleep assay paves the way for future screens aimed at uncovering these processes.

## Acknowledgements

We thank the UCL Fish Facility for animal husbandry, Sumi Lim for genotyping and fish work, Chanpreet Singh for assistance with the tapping assay, Eirinn Mackay for assistance with high speed tracking, David Whitmore for the *per3:luc* assay, Marcus Ghosh for MAP-mapping advice, and Chintan Trivedi, Daniel S. J. Miller and members of the Rihel lab for critical reading of the manuscript. This work was supported by a Sir Henry Wellcome Trust Fellowship (S.R.), a Wellcome Trust PhD Fellowship (O.P.A.), a UCL Excellence Fellowship (J.R.) and a European Research Council Starting Grant (J.R.).

## Author’s contribution

S.R. and J.R. designed and S.R. performed all experiments, except the MAP-mapping, which was performed by O.P.A. under supervision of S.R. S.R. and J.R. wrote the paper with assistance from O.P.A.

## STAR Methods

### EXPERIMENTAL MODEL AND SUBJECT DETAILS

Zebrafish husbandry and experiments were conducted following standard UCL fish facility protocols under project licenses 70/7612 and PA8D4D0E5 awarded to JR from the UK Home Office, according to the UK Animals (Scientific Procedures) Act 1986. Zebrafish AB/TL larvae were studied before the onset of sexual maturation between 5-7 dpf. The age of the animals used is described in the STAR Methods and/or indicated in the figures.

#### Mutant zebrafish

##### galn^int3/int3^ and galn^i8/i8^ mutant

*galn* mutants were generated on a AB/TL background using CRISPR/Cas9 mutagenesis (Hwang et al., 2013) by co-injection of Cas9 mRNA with the sgRNA target sequence 5’-CGGACTCACGAGGACCGAGGA-3’, which is identical in sequence to the boundary of *galn* exon3/intron3. Putative F0 founders harbouring somatic mutations were identified by PCR amplification (primers as below) of a 117 base pair sequence with adaptor arms from tail genomic DNA using a benchtop Illumina sequencing machine (MiSeq). The mutant *galn*^*int3*^ allele harbors a 43 bp deletion spanning the exon3/intron3 boundary causing the loss of the splice donor site in intron3. The *galn*^*int3*^ and wild type alleles were genotyped using the primers 5’-ATGTACTGTCCTCATGGCAAAG-3’ and 5’-AAATGTAGACCTGAGAGCAGC-3’, WT (512 bp) and mutant PCR products (469 bp) were separated by gel electrophoresis on a 2% agarose gel. The mutant i8 allele contains an 8 bp insertion leading to a frame shift and premature stop codon in both *galn* isoforms. The *galn*^*i8*^ and wild type alleles were genotyped as above and wild type and mutant PCR products were separated by gel electrophoresis on a 3% agarose gel. Both *galn* mutant alleles are homozygous fertile and lack any overt morphological or behavioral phenotypes apart from a disorganisation/lack of pigmented stripes due to an impairment of xanthophore development, as melanophores appear unaffected.

### METHOD DETAILS

#### Locomotor activity assay

Behavioral tracking of larval zebrafish was performed as previously described (Prober et al., 2006; Rihel et al., 2010). Zebrafish larvae were raised on a 14:10 light:dark cycle at 28.5°C and at 5 dpf placed into individual wells of a square-well 96-well plate (Whatman) containing 650 µl of standard embryo water (0.3 g/L Instant Ocean, 1 mg/L methylene blue, pH 7.0). Water levels were topped up each morning at the beginning of the light period (9 a.m.) (visible as an additional morning peak in the *galn* mutant behavior traces). Locomotor activity was monitored using an automated videotracking system (Zebrabox, Viewpoint LifeSciences) in a temperature-regulated room (26.5°C) and exposed to a 14:10 white light:dark schedule with constant infrared illumination (Viewpoint Life Sciences). Larval movement was recorded using the Videotrack quantization mode with the following detection parameters: detection threshold, 20; burst, 100; freeze, 3; bin size, 60s, which were determined empirically. At 6 dpf drugs or vehicle alone were added to the individual wells between 10 and 11 a.m. for one hour. Tracking was then paused and larvae were individually placed into 20 cm dish containing embryo water for 5s before being transferred into the well of a fresh 96-well plate corresponding to the larva’s previously tracked well. Tracking was continued until at least the following day. Behavioral tracking of *galn* mutants was performed blinded from heterozygous *galn/+ x galn/+* incrosses. Larvae were genotyped as described above after completion of the experiment and genotypes assigned to wells for data analysis.

The locomotor assay data were analyzed using custom MATLAB (MathWorks) scripts. Any one-minute period of inactivity was defined as one minute of sleep (Prober et al., 2006). Sleep bout length describes the duration of consecutive, uninterrupted minutes of sleep whereas sleep bout number counts the occurrence of such events during a given time period. Average activity represents the total ΔPixel change over time whereas waking activity excludes rest bouts.

#### Inactivity reversibility assay

To test the reversibility of inactivity after drug exposure, larvae were exposed to 10min dark pulses to induce a startle with the first lights-OFF period 20min after drug washout, then every 2.5h until the circadian lights-OFF period (11 p.m.). The response was measured as Δactivity, which was determined for each individual larva by subtracting the mean activity 10min prior to the dark pulse from the mean activity during the 10min dark light period. For pulses 2-4, which on a population level were statistically not significantly different to each other, the average Δactivity response of each larva was calculated.

#### Circadian rhythm assessment

Zebrafish larvae were placed into the videotracking system at 4 dpf and their behaviour monitored over several days in constant light conditions without external perturbation. For qRT-PCR analysis (see below), larvae were raised in constant dark conditions and samples for RNA extraction collected every 6 hours. *per-3* expression was used to compare circadian rhythmicity between wild types and *galn* mutants.

#### *per3* bioluminescent assay

After PTZ exposure, larvae were placed into 96-well plates in water containing 0.5 mM luciferin.Bioluminescence *per3-luciferase* transgenic zebrafish was monitored in constant dark using a Topcount NXT scintillation counter (Packard) at 28°C.

#### Arousal threshold assay

The arousal threshold assay was performed as previously described (Singh et al., 2015). After 1h exposure to 5mM PTZ at 6 dpf, the drugs were washed-out as described above and taps of 14 different intensities were randomly applied over an 8h period (1 p.m. to 9 p.m.). Thirty trials were performed at each stimulus strength, with a 1min interval between trials. The probability of movement independent of the stimulus was calculated by identifying the fraction of larvae for each condition that moved 5s prior to all stimuli delivered. This value was subtracted from the average response fraction value for each tap event. Any movement occurring within 1s of a tap was counted as a response. The curves were analysed using the createFit_v1(x, y) MATLAB function (MathWorks).

#### Sleep deprivation (OMR)

For sleep deprivation using the optomotor response (OMR), 100 larvae were transferred into two clear, square 12 cm petri dishes (Greiner, 688102) in standard embryo water and at 8 p.m. placed above a monitor displaying either a moving (25mm/s) or non-moving grating pattern (control), consisting of black and white vertical stripes. To keep larvae at a constant ambient temperature a fan provided constant air circulation and the water temperature as well as illumination levels were monitored over the course of the experiment. At 10.30 a.m. the following day, 48 larvae of each condition were placed into individual wells of a 96-well plate and their locomotor activity tracked for the next 24h (see above). Exposure of larvae to OMR during the day started at 10.30 a.m. for the duration of 10h. Larvae were then individually tracked for 14h in constant light condictions to assess sleep/wake behaviour.

#### Drug exposure for ISH/IHC/qRT-PCR

Wild type larval zebrafish from the same clutch were raised on a 14:10 light:dark cycle at 28.5°C with lights on at 9 a.m. and off at 11 p.m. At 6 dpf, larvae were transferred to a 6-well plate at a density of up to ~5 fish/mL. At 10:30 a.m. vehicle or drugs were added (concentrations for each experiment are indicated in the figure or figure legend). Larvae were placed in the 28.5°C incubator for 1 hour, after which drugs were washed out and larvae transferred to a new 6-well plate with fresh water. For samples treated for pERK/tERK stainings, larvae were placed into a sieve, which was fitted in each well to facilitate a rapid and consistent fixation across treatment groups and to minimise artifacts in the pERK signal (Randlett et al 2015). After a 4-hour recovery period in the incubator, larvae were quickly fixed by immersing the sieve into a new 6-well plate containing 4% paraformaldehyde (PFA) in phosphate buffered saline (pH 7.3, PBS) + 4% sucrose and left overnight at 4°C.

#### Whole-mount in situ hybridisation (ISH)

Larvae were fixed in PFA with 4% sucrose overnight at 4°C, transferred into PBS the next morning and the brain dissected by removing skin, cartilage and eyes with forceps. ISH was performed following standard protocols using digoxigenin (DIG) −11-UTP-labelled antisense riboprobes targeting the gene transcript of interest (Thisse and Thisse, 2008). Probe hybridization was carried out in hybridization buffer supplemented with 5% dextran sulphate overnight at 65°C. The larvae were then incubated overnight at 4°C with anti-Dig-AP (1:2000 in 5% normal goat serum) and extensively washed before detecting alkaline phosphatase using NBP/BCIP. The antisense *c-fos* riboprobe was transcribed from plasmid after subcloning a 716 bp PCR product generated using primers 5’-CCGATACACTGCAAGCTGAA-3’ and 5’-ATTGCAGGGCTATGGAAGTG-3’ (digestion with BamHI, transcribed with T7 RNA polymerase). The *galn* riboprobe has been previously described (Chen et al., 2017).

Fluorescent ISH was essentially performed as above but probe detection was carried out with an anti-Dig-HRP antibody (1:1000 in 5% NGS). Samples were then developed with the Cy3 TSA® Plus Fluorescence System (Perkin-Elmer, NEL753001KT), larval brains were mounted in 80% glycerol / 1% low melting agarose and imaged using a Leica SP8 confocal microscope with a 25x objective (0.95 NA, water immersion). DAPI was used as a nuclear counterstain. Imaris Image Analysis Software (Bitplanes) was used to count the sum voxel intensity in the selected regions of interest (ROI), which were defined the same size across samples. Mean and median was calculated for larvae processed in the same experiment.

#### Immunohistochemistry (IHC)

Larvae were fixed in 4% PFA with 4% sucrose for 2h (Galanin IHC) or 6h (all other) at room temperature (RT) prior to brain dissection (see above). The samples were washed and pre-incubated with blocking solution in PBS containing 0.25% Triton X-100 (PBS-Tr) with 2% dimethyl sulfoxide (DMSO) supplemented with 2% NGS and for 2h prior to incubation with the primary antibody over night at 4°C. After thorough washing in PBS-Tr, the larvae were incubated with the secondary Alexa antibody (1:500) and DAPI (1 mg/ml stock, 1:1000) overnight at 4°C. Samples were imaged as specified above.

#### Immunohistochemistry (IHC) for pERK/tERK staining

Brains were manually dissected and immunostained as previously described ((Randlett et al., 2015)). Briefly, samples were washed in PBS-Tr three times for 5 min on a shaker, and incubated in 0.05% Trypsin-EDTA for 45 min on ice. After three 5 min washes in PBS-Tr, samples were blocked in PBT + 2% NGS + 1% bovine serum albumin (BSA) + 1% DMSO at RT on a shaker for at least 1 hour. Primary antibody incubations were performed overnight at 4°C on a shaker in PBS-Tr + 1% BSA + 1% DMSO using mouse anti-tERK (1:500) and rabbit anti-pERK (1:500). Samples were washed three times for 15 min in PBS-Tr at RT and secondary antibody incubations were performed overnight at 4°C on a shaker in PBS-Tr + 1% BSA + 1% DMSO using Alexa Fluor 488 goat anti-rabbit (1:200) and Alexa Fluor 568 goat anti-mouse (1:200). Finally, samples were washed three times for 15 min in PBS-Tr at RT and then left in PBS-Tr rotating at 4°C until mounting.

To allow registration of the signal obtained by fluorescent ISH against *galn*, brains were immunostained as described above using mouse anti-tERK (1:500) as primary antibody and Alexa Fluor 568 goat anti-mouse (1:200) as secondary antibody.

#### Image acquisition, Z-brain registration and MAP-mapping

Stained samples were imaged in a custom-built 2-photon microscope (Bruker), using a 20x water-immersion objective (1.0 NA, OLYMPUS) and Prairie View acquisition software. Brains were mounted dorsal-side up in 1.5% low melting point agarose and full-brain stacks were acquired at a voxel size of 0.584/0.584/2 µm (x/y/z) with a laser wavelength of 800 nm and a laser intensity of 40-80 mW. Brains that underwent fluorescent ISH against *galn* together with IHC against tERK were imaged using a Leica SP8 confocal microscope with a 25x objective (Olympus, 0.95 NA, water immersion at a voxel size of 0.606/0.606/1.5 µm (x/y/z).

Analysis was performed using MATLAB (MathWorks) and Fiji. Non-rigid image registration was performed using the Computational Morphometry Toolkit (CMTK, http://www.nitrc.org/projects/cmtk/) as previously described (Randlett, 2015). tERK staining was used to register the samples to the Z-brain atlas reference brain (Randlett et al., 2015) and then used to align pERK stainings, as well as the galanin FISH labeling. Calculation of MAP-maps was done using the "MakeTheMAPMap.m" Matlab function as previously described (Randlett et al., 2015). Maximum-intensity projections of MAP-maps were obtained after resizing the raw MAP-maps to 621×1406 pixels and 138 slices in Fiji. Lateral views of maximum-intensity projections of MAP-maps were obtained by resizing the stacks to 276 slices depth and reslicing them in Fiji (1 micron output spacing, start at left, rotate 90 degrees, avoid interpolation).

To further analyse and quantify the MAP-map signals as well as the signal from the registered *galn* ISH, we compared their distribution with the regions and cell type labels in the Z-Brain atlas using the “ZBrainAnalysisOfMAPMaps.m” MATLAB function as previously described (Randlett, 2015) (Supplementary Table 1).

#### Quantitative real-time PCR (qRT-PCR)

For qRT-PCR analysis, 30-40 whole larvae in minimal medium were snap frozen in liquid nitrogen. RNA was isolated using TRIzol™ Reagent (Invitrogen, 15596026) and treated with DNaseI (NEB, M0303). 1 µg of RNA was reverse transcribed with AffinityScript (Agilent, 600559) the resulting cDNA was diluted 1:20. Expression levels were measured using GoTaq® qPCR Master Mix (Promega, A6002) in technical triplicates on a BioRad iCycler. For each primer set, efficiency and specificity was first determined using standard curves of diluted cDNAs and melting curve analysis. Expression of *galn* and *fosab* (*c-fos)* was normalized to expression of *ef1a1l1* and relative expression levels were calculated using the ΔΔCt method. Statistical tests for differences are indicated in the figure legends. Primers for amplification: *galn*, 5’-AAGGATACTCCCAGTGCAAGG-3’ and 5’-CTTTCCTGCCAGTCCGTGTT-3’; *fosab*, 5’-GTGCAGCACGGCTTCACCGA-3’ and 5’-TTGAGCTGCGCCGTTGGAGG-3’ (Ellis et al., 2012); *ef1a1l1*, 5’-TGCTGTGCGTGACATGAGGCAG-3’ and 5’-CCGCAACCTTTGGAACGGTGT-3’.

#### Wide field Calcium (Ca^2+^) imaging

Wide field microscopy imaging of whole brain Ca^2+^ signals was performed as previously described with the following modifications (Turrini et al., 2017). Neuronal activity was measured in 6 dpf larvae expressing the genetically-encoded calcium indicator GCaMP6s under the pan-neuronal promoter elav3 (Tg(elavl3:GCaMP6s)). Larvae were mounted in a drop of 3% low melting agarose in standard embryo medium in a 35 mm petri dish (Falcon, 353001) and the dish filled with standard embryo medium. Imaging was performed on an upright MVX10 MacroView microscope using an MC PLAPO 1x objective (both OLYMPUS) with a mercury lamp for fluorescent excitation (OLYMPUS, U-HGLGPS). The fluorescence signal was recorded with XM10 OLYMPUS camera at 5.5 frames per second (fps). Baseline was recorded twice over a 5 min period prior to addition of the drug or vehicle control. After exposure to DMSO, 10 mM PTZ or 5 µM aconitine, fluorescence was immediately recorded for 5 min every 5 min over a 65 min period (total of 7×5 min recordings). The GCaMP6s fluorescence signal was measured using ImageJ by manually selecting the brain as the region of interest (ROI). After background subtraction, ΔF/F was calculated using the mean pre-drug baseline fluorescence. For global comparison between conditions, the fluorescence per min (normalised to pre-treatment baseline) was calculated for each of the 5 min recordings and the average taken over the 7 repeats.

### QUANTIFICATION AND STATISTICAL ANALYSIS

For behavioral experiments, after testing for deviations from normality, Kruskal-Wallis multiple comparison followed by Dunn-Sidak post-hoc test was used to assess statistical significance. Experimenters are blinded to genotype.

For *galn* and *c-fos* qPCR to measure induction of expression, one-way ANOVA with post hoc Dunnett’s test for comparisons to controls was used. Quantification of *galn* expression time course post PTZ and return to baseline the following day was tested for significance using one-way ANOVA for pairwise comparisons of the mean of each sample to its timed control with post hoc Bonferroni correction for multiple comparisons. Differences in *galn* expression using fluorescent ISH were assessed by Kruskal-Wallis multiple comparisons followed by Dunn-Sidak post-hoc test. Statistical analysis of *galn* mutants in response to OMR SD was assed by two-way ANOVA (genotype x drug condition) followed by a Tukey-Kramer post-hoc test. MATLAB and PRISM were used for statistical analyses. Statistical tests (where different from above description) and number of animals (n) used are stated in each figure or figure legend.

### DATA AND SOFTWARE AVAILABILITY

Custom MATLAB code used for zebrafish behavioral analysis is available upon request.

---

**Figure S1.**
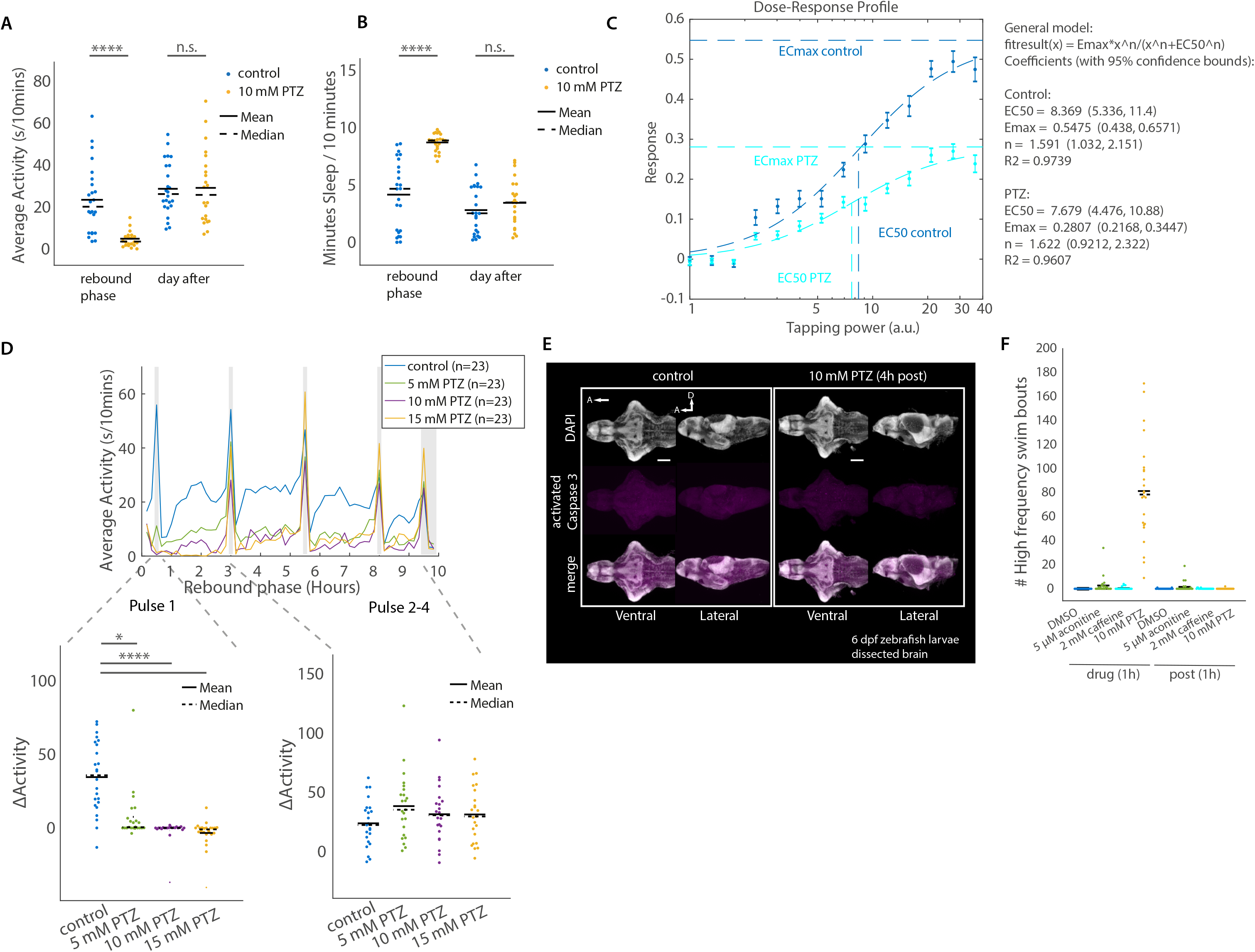
Related to Figure 1. PTZ exposure is followed by rebound sleep. (A and B) Quantification of average activity (A) and average sleep (B) during the rebound phase compared to the following day shows a full behavioral recovery of PTZ treated larvae. ****p<0.0001, n.s., p>0.05 (C) Stimulus response curves for 10 mM PTZ treated larvae (n=34) and controls (n=44) during rebound phase. Each data point represents mean response fraction ±SEM to acoustic taps of varying strength (arbitrary units, a.u., see Extended Methods). The EC_50_ and EC_max_ are indicated. (D) Dark pulses during the rebound phase reverse larval inactivity in both PTZ-treated and untreated larvae after an initial refractory period (Pulse 1). Each trace plots the 10 minute average of n=23 larvae from a representative experiment. Grey bars indicate the timing of 10-minute dark pulses during the rebound phase. The change in activity (ΔActivity), calculated as the change in activity per 10 minutes of the dark pulse minus the preceding 10 minute activity for individual larvae is shown below for the first dark pulse (left) and the average of pulses 2-4 combined (right). *p<0.05, ****p<0.0001 (Pulse 1), p=0.34 (Pulse 2-4) (E) Caspase3 staining on dissected larval brains (magenta), as an indicator of apoptosis, shows no increase in cell death in the brains of 6 dpf larvae treated with PTZ for 1h and collected 4h into the rebound phase, as compared to the control group. DAPI nuclear staining is labelling cell nuclei (grey). (Scale bar: 100 microns. A, Anterior; D, Dorsal. (F) Quantification of high amplitude swim bouts recorded during one hour of drug application (drug (1h), doses indicated) and during one hour directly after wash out. n=24 for each condition.

**Figure S2.**
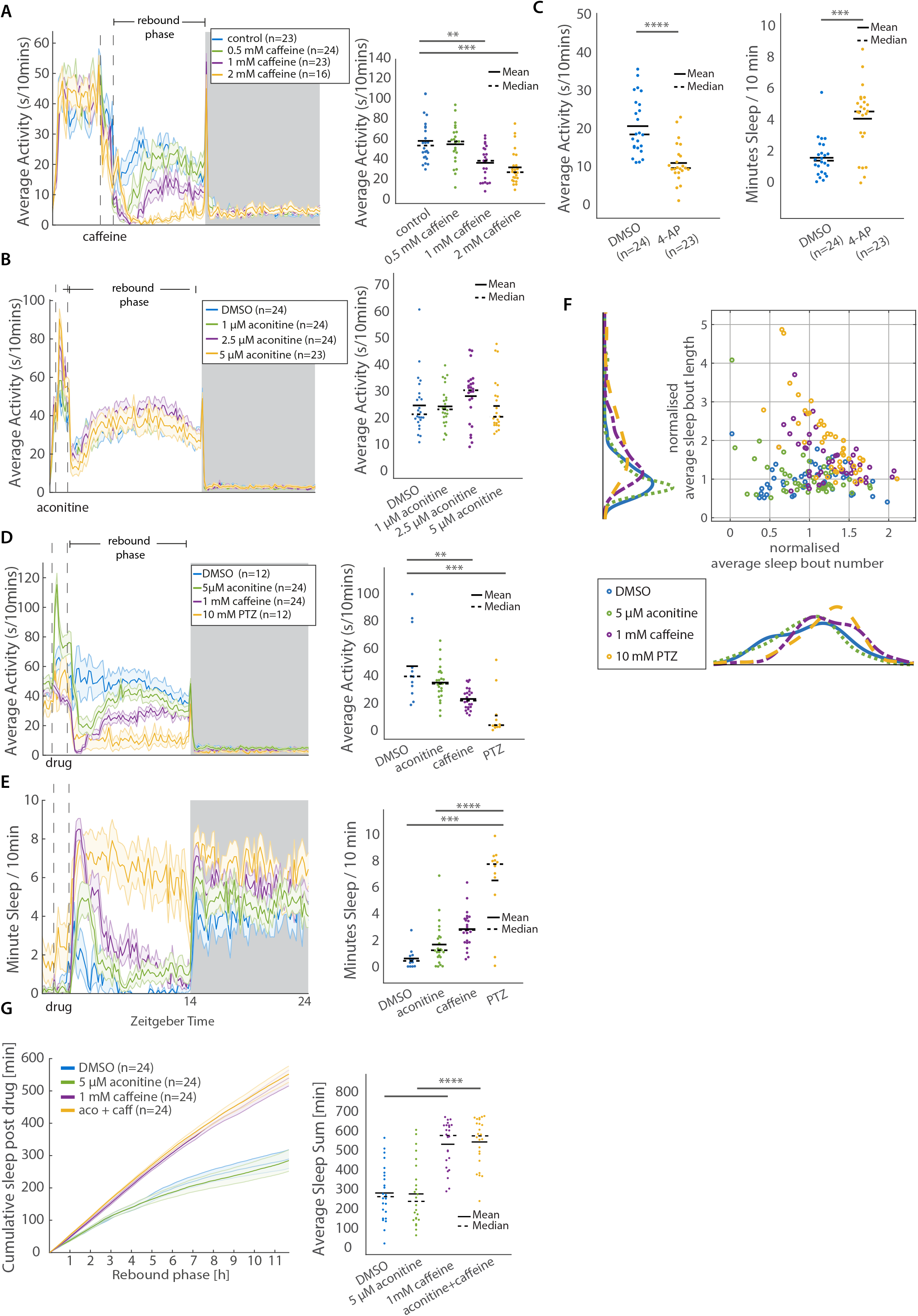
Related to Figure 2. Caffeine, but not aconitine, induces rebound sleep. (A) Locomotor behavior is reduced after 1h treatment with three doses of caffeine (rebound phase). The dashed line indicates timing of caffeine treatment, while the light and dark boxes indicate lights ON and OFF periods, respectively. The ribbon along each trace depicts ±SEM. Average data for individual larvae during the rebound phase is shown to the right of each behavior trace. **p≤0.01, ***p≤0.001, ****≤0.0001 (B) Locomotor behavior remains unchanged after 1h treatment with three doses of aconitine. The dashed line indicates timing of aconitine treatment, while the light and dark boxes indicate lights on and off periods, respectively. The ribbon along each trace depicts ±SEM. Average data for individual larvae during the rebound phase is shown to the right. n.s. p>0.05 (C) Locomotor behavior is reduced and sleep increased after 1h treatment with 400 µM 4-AP. Average data for individual larvae is shown. ***p≤0.001, ****≤0.0001 (D and E) Locomotor (D) and sleep (E) behavior after 1h treatment with aconitine, caffeine, or PTZ. The dashed line indicates timing of drug treatment, while the light and dark boxes indicate lights on and off periods, respectively. The ribbon along each trace depicts ±SEM. Average data for individual larvae during rebound phase is shown adjacent to behavior trace. **p≤0.01, ***p≤0.001, ****≤0.0001 (F) Scatter plot showing average sleep bout length and number during the rebound phase for individual larvae for two independent experiments, normalised to the population mean of the control group. Along the axes, histograms show the distributions for average sleep bout number (x-axis) and mean sleep bout length (y-axis) for each condition. (G) Cumulative sleep after 1h exposure to indicated drugs (rebound phase). The ribbons represent ±SEM. Total accumulated sleep for individual larvae is shown to the right. ****p≤0.0001

**Figure S3.**
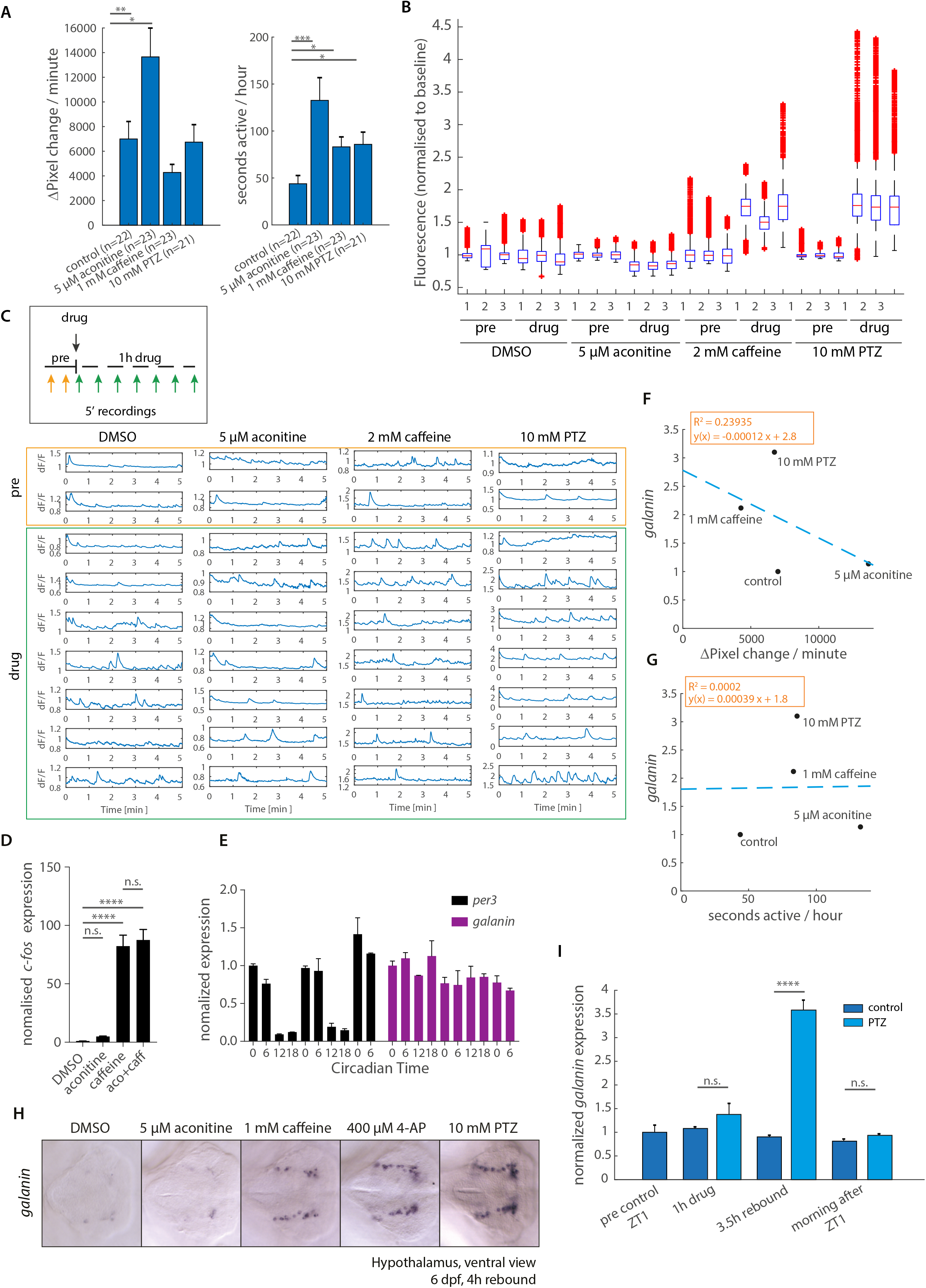
Related to Figure 3 and 4. Global measures of neuronal activity correlate with rebound sleep amount. (A) Bar charts showing mean movement measured over 1h as Δpixels (Left) and active seconds per hour (Right) in response to treatment with the indicated pharmacological compounds. *p<0.05, **p≤0.01, ***p≤0.001 (B) Whole brain fluorescent Ca^2+^ signals of individual frames before (pre) (mean of 2 x 5min recordings) and after addition of pharmacological compounds (drug) (mean of 7 x 5min recordings over an hour) are shown for three individual larval brains per condition. Fluorescent signals were imaged at 5.5 frames per second (fps) and signals normalised to the average baseline recorded pre drug. (C) Traces of representative, individual whole brain fluorescent Ca^2+^ signals before (orange box) and during (green box) addition of the indicated drugs. Imaging was performed at 5.5 fps over 5min time windows. All traces were normalised to the average pre-recording baseline. Note the changes in y-axis scale across samples. (D) qRT-PCR analysis of *c-fos* in response to 5 µM aconitine, 1 mM caffeine or a combination. One-way ANOVA followed by Tukeys multiple comparison test. ****p≤0.0001 (E) qRT-PCR analysis of the circadian gene *per3* and *galn* over 2.5 days in constant dark conditions. (F and G) A weak negative correlation (F) or no correlation (G) between behavioural activity measures from A (x-axis) and *galn* expression (y-axis) as measured by RT-qPCR. The dashed line depicts a linear regression curve (R^2^=0.24 and R^2^=0.0002). (H) *galn* ISH on dissected brains collected 4h into rebound phase after 1h exposure to indicated drugs shows increased *galn* expression after sleep rebound inducing drugs but not after aconitine. Ventral view of larval zebrafish hypothalamus. (I) qRT-PCR analysis comparing *galn* expression levels between controls and 10 mM PTZ-treated samples at four time points: pre experiment start, directly after 1h drug treatment, 3.5h into the rebound phase (*galn* up), and the morning of the following day (*galn* returned to baseline). All data points normalized to pre-drug controls. n.s. p>0.05, ****p≤0.0001, one-way ANOVA with post hoc Bonferroni correction for comparison to timed controls.

**Figure S4.**
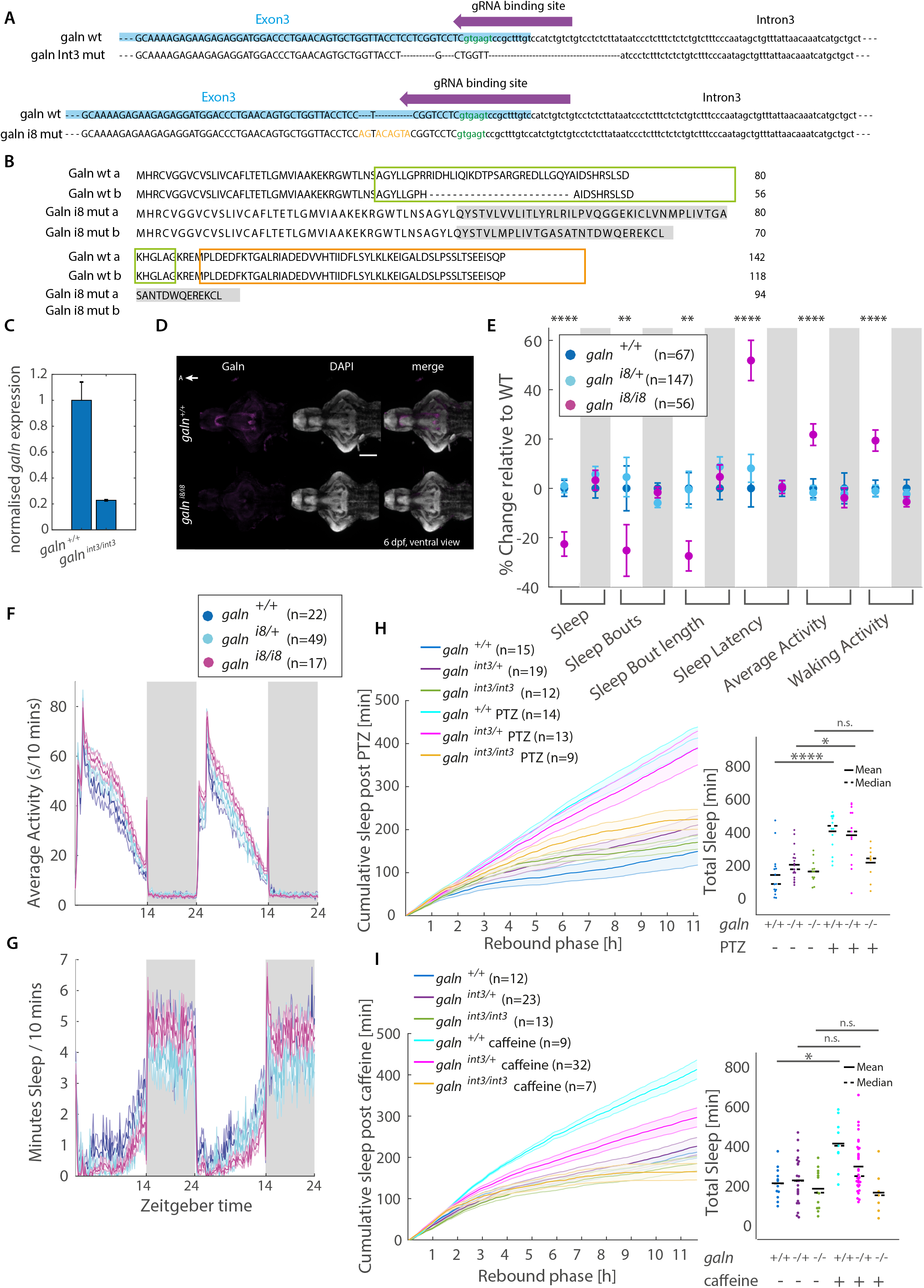
Related to Figure 5. Galanin is required for neural activity induced rebound sleep. (A) The *galn* gene locus was targeted by Crispr/Cas9 at the boundary between exon 3 (blue upper case) and intron3 (black lower case) to generate two mutant alleles. The gRNA binding site for directed Crispr/Cas9 targeting is indicated with a purple arrow. The *galn*^*int3*^ allele carries a mutation leading to the loss of splice donor site (green) (top), whereas the *galn*^*i8*^ allele has an 8bp insertion (orange). Both are predicted to encode loss of function mutations. (B) The amino acid sequence of wild type and *galn*^*i8*^ mutant Galn preproprotein (isoforms a and b). The Galn^i8^ predicted protein is missing most of the mature Galn peptide (green box) and Galn Message Associated Peptide (orange box). Grey shading highlights altered amino acids in the predicted Galn^i8^ mutant. (C) *galn* mRNA expression is down regulated in *galn*^*int3/int3*^ mutants compared to wild type siblings (qRT-PCR analysis), consistent with nonsense mediated decay. ***p≤0.001, one-way ANOVA. (D) Immunofluorescence staining detects no Galn protein in 6 dpf *galn*^*i8/i8*^ mutant brains compared to wild type siblings (magenta). Nuclei are stained with DAPI (grey). Scale bar: 100 microns. A, anterior (E) Behavioral profile of *galn*^*i8/i8*^ mutants shown as the mean % change relative to wild type siblings for indicated features of larval sleep architecture for day (white background) and night (grey background). Error bars indicate ±SEM from three independent, blinded experiments. *p≤ 0.05, **p≤0.01, ***p≤ 0.001, ****p≤ 0.0001. (F and G) Locomotor (F) and sleep (G) behavior traces for a representative experiment comparing *galn*^*i8/i8*^, *galn*^*+/i8*^, and *galn*^*+/+*^ siblings. White and grey areas indicate lights-ON and lights-OFF periods, respectively. The ribbons depict ±SEM. (H and I) Cumulative sleep during rebound phase after exposure to 5 mM (H) or 1 mM (I) caffeine. The ribbons depict ±SEM. Quantification of total average cumulative sleep during the rebound phase at the end of the circadian light period is shown next to behavior traces. *p≤0.05, *p≤ 0.05, ****p≤0.0001

**Figure S5.**
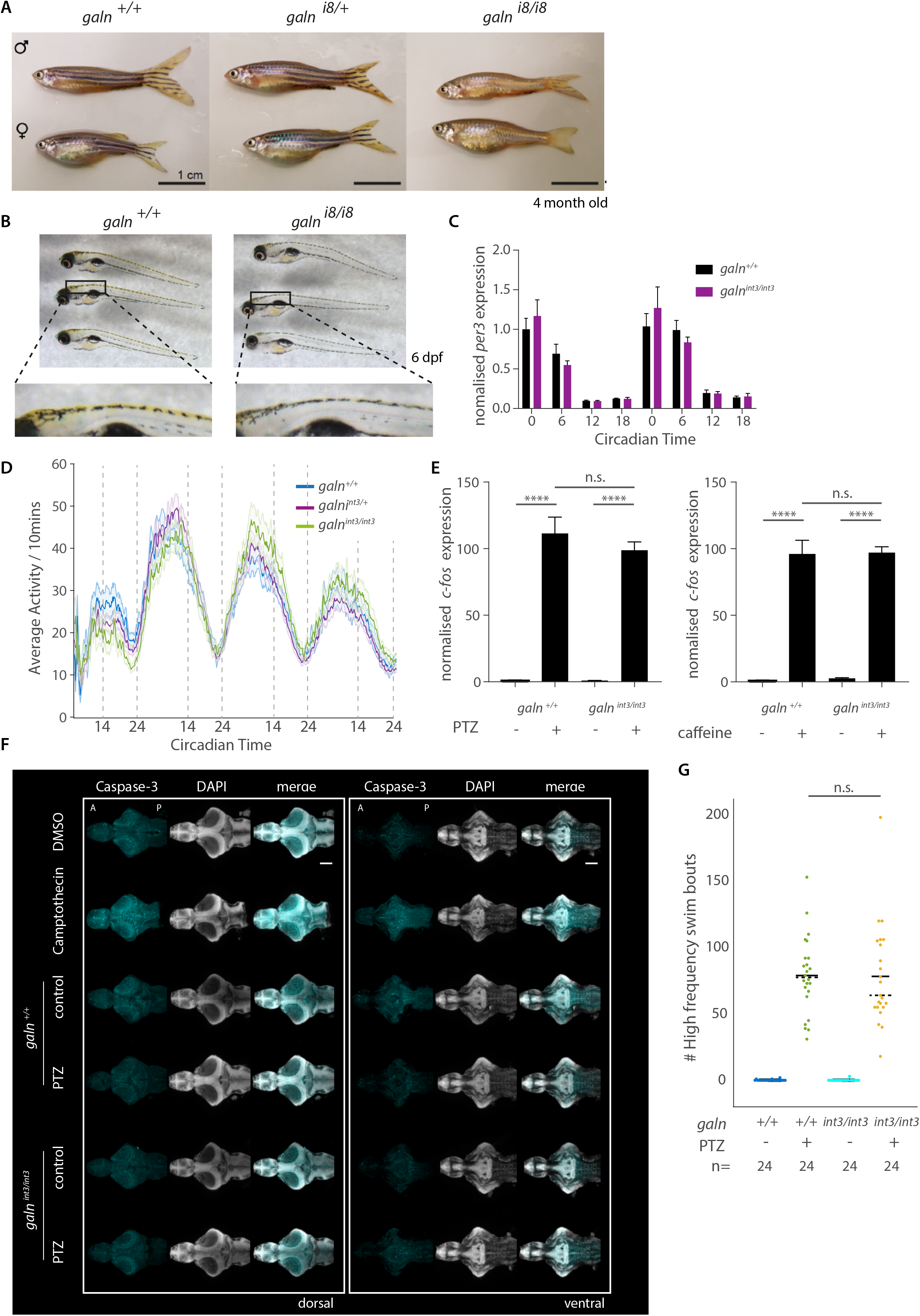
Related to Figure 5 - Galanin mutants characterisation. (A and B) Pigmentation phenotype in *galn*^*i8/i8*^ mutant adults (A) and 6 dpf larva (B). Note that melanophores seem to be unaffected but yellow-pigmented xanthophores are lost (highlighted in (B), lower panels). The same phenotype is observed in *galn*^*int3/int3*^ mutants. (C) qRT-PCR analysis of the circadian gene *per3* shows no difference in expression amplitude or phase in *galn*^*int3/int3*^ mutants compared to wild type. (D) Activity behaviour trace of *galn*^*+/+*^, *galn*^*int3/+*^ and *galn*^*int3/int3*^ larvae in constant light conditions. Dashed lines indicate subjective day/night transitions. (G) qRT-PCR analysis of c-fos expression in *galn*^*int3/int3*^ mutants in response to 10 mM PTZ and 2 mM caffeine. Two-way ANOVA (gene x drug) followed by Tukey-Kramer post-hoc test. ****p≤ 0.0001 (F) Caspase3 staining on dissected larval brains (cyan), as an indicator of apoptosis, shows no increase in cell death in the brains of 6 dpf *galn*^*+/+*^ or *galn*^*int3/int3*^ mutant larvae treated with 10 mM PTZ for 1h and collected 4h into the rebound phase. Larvae were exposed to the topoisomerase inhibitor Camptothecin (2 µM) as a positive control for apoptisis. DAPI nuclear staining is labelling cell nuclei (grey). (Scale bar: 100 microns. A, Anterior; D, Dorsal. (G) Quantification of high burst swim bouts of *galn*^*+/+*^ or *galn*^*int3/int3*^ mutant larvae recorded during one hour 5 mM PTZ application and during one hour directly after wash out. n.s., p>0.05

**Figure S6.**
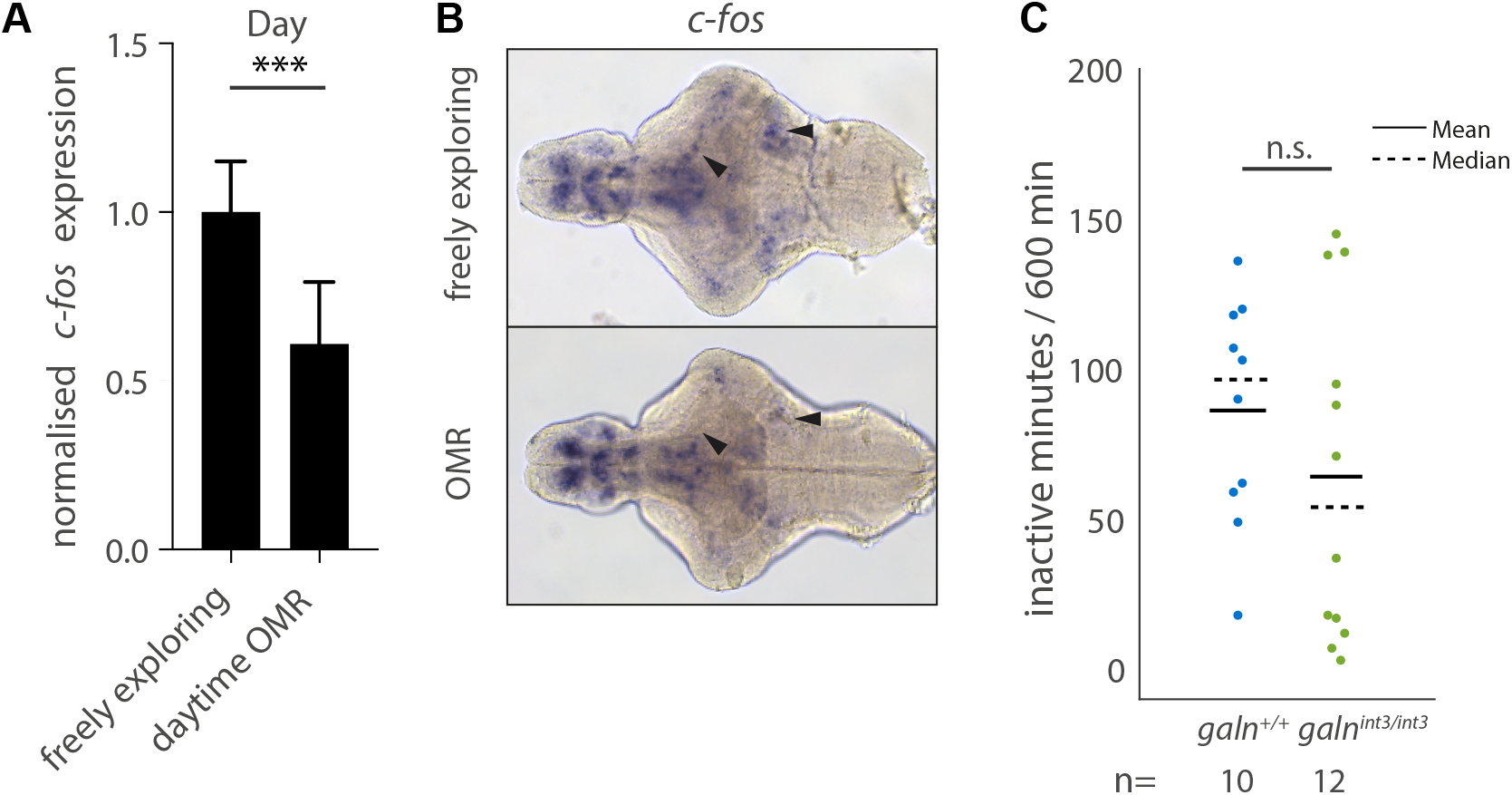
Related to Figure 6 – OMR sleep deprivation. (A) RT-qPCR analysis measuring *c-fos* after day time OMR exposure compared to freely exploring larvae. ***p≤ 0.001, one-way ANOVA. (B) ISH hybridisation visualising the spatial expression of *c-fos* expression after free exploration or exposure to day time OMR (black arrows highlight areas of differential expression). (C) Analysis of time spend inactive during night time OMR SD in *galn*^*+/+*^ and *galn*^*int3/int3*^. Each dot represents a single larva. n.s. p>0.05, Mann–Whitney U test.

**Supplementary Table 1**

Table shows a list of anatomical regions and cell type labels containing functional signals from the MAP-maps (Fig. 5 and S5, see Extended Methods) and the registered *galn* ISH after comparing the signal distribution with the Z-Brain atlas.

**Supplementary Movie 1-3**

Videos show whole brain Ca^2+^ imaging of representative Tg(elav3:H2B-GCaMP6s) larval brains recorded over 5 min at 5.5 fps and replayed at 500 fps. Movie 1 – 5 µM aconitine; Movie 2 – 1 mM caffeine; Movie 3 – 10 mM PTZ.

